# The role of climatic anomalies in agricultural instability and settlement abandonment in northeastern Germany during the Migration Period

**DOI:** 10.1101/2025.10.08.681122

**Authors:** Khadijeh Alinezhad, Rongwei Geng, Evelien J. C. van Dijk, Wiebke Kirleis, Mara Weinelt

## Abstract

The Late Antique Migration Period (LAMP; 300–700 CE) in northeastern Germany was marked by significant socio-environmental transformations, including the decline of human activity, settlement abandonment, and a reduction in cereal cultivation, particularly rye (*Secale cereale*). While these changes are well-documented in pollen and archaeological records, their underlying drivers—whether socio-political instability, agricultural crises, or environmental pressures—remain debated. Here, we synthesize multiproxy evidence, including high-resolution pollen data, climate reconstructions, and archaeobotanical data to explore the interplay between climatic fluctuations, agricultural practices, and human mobility during LAMP.

Although previous studies have examined socio-environmental changes during the Migration Period using archaeological and palynological evidence, this study is the first to integrate high-resolution pollen records, archaeobotanical data, and climate model simulations specific to northeastern Germany to systematically evaluate the role of climatic anomalies in shaping agricultural and demographic patterns. We focus on the Dark Age Cold Period (DACP; 350–700 CE) and the Late Antique Little Ice Age (LALIA; 536–660 CE), which brought cold and wet conditions to northern Europe, likely disrupting agricultural productivity and fostering the spread of *Claviceps purpurea* (ergot), a toxic fungus affecting cereal crops, particularly rye. Our findings suggest that these climatic anomalies created favorable conditions for ergot outbreaks, leading to agricultural instability and reduced food security. This, combined with socio-political pressures, may have contributed to the observed decline in human activity and settlement abandonment.

By integrating palynological, climatic, and archaeobotanical data, this study highlights the complex interactions between environmental stressors and human responses during the LAMP. We suggest that the climatic anomalies of the DACP and LALIA, coupled with the vulnerability of cereal crops to ergot infestation, influenced agricultural and demographic shifts in northeastern Germany. This review not only refines our understanding of the DACP and the LALIA but also demonstrates how climatic stressors and natural hazards can amplify socio-political vulnerabilities in past societies (or pre-industrial societies).

## 1. Introduction

Pollen records and archaeological evidence reveal that during the Late Antique Migration Period (LAMP; 300–700 CE), northeastern Germany experienced significant socio-environmental changes, marked by declining human activity, reduced cereal cultivation—particularly rye (*Secale cereale*)—and subsequent forest regeneration (Clarke 1985 p 40; Ejstrud et al 2008; cf. Volkmann 2014, p 137–138; Jahns et al 2013a, 2013b, 2018; Alinezhad et al 2024). While these patterns indicate a clear trajectory of settlement abandonment and agricultural decline, the underlying drivers of these socio-environmental shifts remain poorly understood. Current debates focus on whether these shifts were primarily driven by socio-political instability, agricultural crises (e.g., crop disease), external pressures (e.g., environmental change due to climatic cooling), or interactions between these factors (Büntgen et al 2011, 2016; Toohey et al 2016; Schneeweiß 2023). Given these uncertainties, a multi-proxy approach is necessary to disentangle climatic and anthropogenic influences.

Northeastern Germany was selected for this study due to its suitability for investigating long-term socio-environmental change, given its location near major river corridors such as the Dosse and Havel, which historically influenced settlement and agricultural patterns. Its transitional climate, between Atlantic and continental regimes, also makes it particularly sensitive to hydroclimatic variability. Furthermore, the region benefits from a rich archaeological record and several high-resolution palaeoenvironmental studies (e.g., Jahns et al 2013a; 2013b; 2018), which offer a strong framework for comparative analysis. Therefore, to answer the question that how climatic anomalies during the LAMP may have influenced agricultural productivity and contributed to the observed decline in human activity we use a multiproxy approach centered on original data plus published datasets to examine the underlying causes of these socio-environmental transformations in northeastern Germany during the LAMP. We present the first integrative study to systematically examine how climatic anomalies contributed to agricultural instability and demographic change during the Late Antique Migration Period, by combining high-resolution palynological data, regional archaeobotanical records, and local-to-regional scale proxy-based climate reconstructions with continental climate model simulations. In doing so, we advance previous research by directly linking climatic anomalies with agricultural instability and demographic change during the Migration Period. First, we use previously published high-resolution pollen records from lake Kleiner Tornowsee (KTO) to show the documented shifts in vegetation, land use, and human activity during the LAMP. Second, using pollen-based climate reconstructions from lake KTO and regional datasets, combined with continental climate model simulations, we assess climatic fluctuations during this period and evaluate their potential influence on the socio-environmental changes during this period. Third, we include archaeobotanical data from domestic sites to provide insights into potential shifts in crop cultivation, including the presence of different cereal species (e.g., rye, wheat, barley) and their abundances. Thus, the present approach overcomes the limitations of earlier studies that relied on single lines of evidence by enabling a more comprehensive understanding of the interactions between agriculture, climate, and human mobility. Notably, while pollen and archaeobotanical data have previously been used to investigate land-use changes, this study incorporates climate model simulations for the region for the first time, thereby offering a novel perspective on how climatic anomalies may have contributed to agricultural instability and demographic decline.

Therefore, by integrating these datasets we evaluate the interplay between human mobility, agricultural change, and environmental conditions in northeastern Germany during the LAMP, which has not been done in this way before.

### 1.1 The Late Antique Migration Period (LAMP)

Between 300 and 700 CE, a period often referred to as the LAMP, Europe underwent significant environmental, social, agricultural, economic, and political changes. This period was characterized by widespread socio-environmental disruptions and human migrations that took place across many European regions (Büntgen et al 2011; Toohey et al 2016; Bajard et al 2022). However, the extent and nature of these transformations, as well as their underlying causes and triggers, remain insufficiently understood.

A key aspect of these transformations in Europe was the major demographic shifts that occurred during this time, which can be divided into two distinct phases: the 3^rd^ to 5^th^ centuries CE and the 5^th^ to 9^th^ centuries CE (Dreßler et al 2006; Meier 2019; Rubini et al 2022; Czerwiński et al 2022). The earlier phase (300-500 CE) saw the migration of Germanic groups (e.g., Goths, Angles, Saxons, Jutes, Vandals) and of warrior groups from northern and eastern Europe into regions such as modern France, the British Isles, the Italian Peninsula, the Iberian Peninsula, Anatolia, and the Maghreb. In the later phase (500– 900 CE), Slavic groups emerged in central and eastern Europe (Dreßler et al 2006; recently for Eastern Europe: Vyazov et al 2024). In northeastern Germany this broader pattern of transformation is reflected in two distinct demographic phases. The first, from the late 4^th^ to mid-6^th^ century CE is characterized by a gradual decline of settlements, which is archaeologically associated with the migration of Germanic groups toward the Roman provinces, particularly Britain (Volkmann 2014, p 137–138). The second phase beginning in the late 6^th^ century and extending into the 7^th^ –9^th^ centuries CE corresponds with the arrival of Slavic groups in central and eastern Europe including northeastern Germany (Dreßler et al 2006; Vyazov et al 2024).

### 1.2 Climatic background

The LAMP period in Europe was marked by two distinct climatic events: the Roman Warm Period (RWP; 300 BCE to 350 CE) and the Dark Age Cold Period (DACP; 350–700 CE) (Büntgen et al 2011, 2016; Riechelmann and Gouw-Bouman 2019; Gouw-Bouman 2023; McKay et al 2024). The DACP, temporally overlapping with the LAMP, is characterized by severe climatic fluctuations, including pronounced cooling and hydroclimatic changes in the Northern Hemisphere; however, recent studies suggest that these Late Holocene changes were regionally rather than globally coherent climate epochs (Neukom et al 2019). For Europe, this period displays consistent patterns of cold and wet conditions (table 1), that were particularly pronounced in northern and western regions. These conditions were likely driven by a prolonged negative phase of the North Atlantic Oscillation (NAO), resulting from sea-level pressure differences between the Azores High and the Icelandic Low. In contrast regions around the Mediterranean and the China/Tibetan Plateau experienced drier conditions during this time, highlighting the global variability of climatic impacts (Helama et al 2017).

**Table 1.** Characteristics of the Dark Age Cold Period (DACP, 350–700 CE) and the Late Antique Little Ice Age (LALIA, 536 to 660 CE) in 19 palaeoclimate papers, with proxy type (Multi-proxy, Sedimentary, Pollen, Dendrology, Other), geographical regions (NWE – NW Europe, ALP – Alps, EAE – East Europe), and the actual proxy indications.

| Reference | Proxy | Region | Climatic Period | Climatic Impact |
| --- | --- | --- | --- | --- |
| Bjune et al. 2009 | Pollen | NWE | DACP | Cold |
| Seppä et al. 2009 | Pollen | NWE | DACP | Cold |
| Gouw-Bouman 2024 | Pollen | NWE | DACP | Woodland regeneration, wet, cold |
| Björck and Clemmensen 2004 | Sedimentary | NWE | DACP | Cold, wet, enhanced storminess |
| Clemmensen et al. 2009 | Sedimentary | NWE | DACP | Cold, wet |
| Larsen et al. 2011 | Sedimentary | NWE | DACP | Cold |
| Dong et al. 2011 | Multi-proxy | NWE | DACP | Cold, storminess |
| Reimann et al. 2011 | Multi-proxy | NWE | DACP | Cold, storminess |
| Stacke et al. 2014 | Multi-proxy | EAE | DACP | Increased fluvial activity |
| Swindles et al. 2013 | Multi-proxy | NWE | DACP | Wet |
| Cristea et al. 2014 | Other | EAE | DACP | Cold, glacial expansion |
| Jakab et al. 2009 | Other | EAE | DACP | Cold |
| Orme et al. 2015 | Other | NWE | DACP | Storminess |
| Rørvik et al. 2009 | Other | NWE | DACP | Cold |
| Helama et al. 2009 | Other | NWE | DACP | Wet |
| McDermott et al. 2001 | Speleothem | NWE | DACP | Cold |
| Jansma 2020 | Dendrology | NWE | DACP | Flooding, wet |
| Helama et al. 2021 | Tree-ring isotope | NWE | DACP | Cold |
| Gouw-Bouman et al. 2019 | Chironomid | NWE | DACP | Cold |
| Büntgen et al. 2016 | Dendrology | ALP | LALIA | Cold |
| Helama et al. 2018 | Tree-ring isotope | NWE | LALIA | Cold |
| Waltgenbach et al. 2021 | Multi-proxy / Speleothem | NWE | LALIA | Cold, wet |

Table 1. Characteristics of the Dark Age Cold Period (DACP, 350–700 CE) and the Late Antique Little Ice Age (LALIA, 536 to 660 CE) in 19 palaeoclimate papers, with proxy type (Multi-proxy, Sedimentary, Pollen, Dendrology, Other), geographical regions (NWE – NW Europe, ALP – Alps, EAE – East ...
| Reference | Proxy | Region | Climatic Period | Climatic Impact |
| --- | --- | --- | --- | --- |
| Bjune et al. 2009 | Pollen | NWE | DACP | Cold |
| Seppä et al. 2009 | Pollen | NWE | DACP | Cold |
| Gouw-Bouman 2024 | Pollen | NWE | DACP | Woodland regeneration, wet, cold |
| Björck and Clemmensen 2004 | Sedimentary | NWE | DACP | Cold, wet, enhanced storminess |
| Clemmensen et al. 2009 | Sedimentary | NWE | DACP | Cold, wet |
| Larsen et al. 2011 | Sedimentary | NWE | DACP | Cold |
| Dong et al. 2011 | Multi-proxy | NWE | DACP | Cold, storminess |
| Reimann et al. 2011 | Multi-proxy | NWE | DACP | Cold, storminess |
| Stacke et al. 2014 | Multi-proxy | EAE | DACP | Increased fluvial activity |
| Swindles et al. 2013 | Multi-proxy | NWE | DACP | Wet |
| Cristea et al. 2014 | Other | EAE | DACP | Cold, glacial expansion |
| Jakab et al. 2009 | Other | EAE | DACP | Cold |
| Orme et al. 2015 | Other | NWE | DACP | Storminess |
| Rørvik et al. 2009 | Other | NWE | DACP | Cold |
| Helama et al. 2009 | Other | NWE | DACP | Wet |
| McDermott et al. 2001 | Speleothem | NWE | DACP | Cold |
| Jansma 2020 | Dendrology | NWE | DACP | Flooding, wet |
| Helama et al. 2021 | Tree-ring isotope | NWE | DACP | Cold |
| Gouw-Bouman et al. 2019 | Chironomid | NWE | DACP | Cold |
| Büntgen et al. 2016 | Dendrology | ALP | LALIA | Cold |
| Helama et al. 2018 | Tree-ring isotope | NWE | LALIA | Cold |
| Waltgenbach et al. 2021 | Multi-proxy / Speleothem | NWE | LALIA | Cold, wet |

The cold and wet conditions of the DACP were further intensified by the Late Antique Little Ice Age (LALIA), a period from 536 to 660 CE characterized by significant temperature drops and disruptive climate and highly unstable weather patterns, triggered by a series of major volcanic eruptions around 536 CE and reduced solar irradiance (Stothers 1984; McCormick et al 2012; Büntgen et al 2016; McKay et al 2024). During the LALIA, remarked climatic and environmental hazards occurred across Europe. These included abrupt climate shifts, which led to interconnected effects such as famine, disease outbreaks, human migrations, and the decline and even fall of civilizations (Baillie 1994; Larsen et al 2008; Büntgen et al 2011; Büntgen et al 2016; Toohey et al 2018; Schneeweiß 2023). These climatic and environmental hazards led to widespread economic and agricultural instability, as temperature drops during the growing season negatively impacted crop yields (Stamnes 2016; Toohey et al 2018; van Dijk et al 2024).

### 1.3 Agriculture

In Central Europe, cereal cultivation had been firmly established as a primary agricultural practice since the Neolithic with a rather small set of cultivars that was widened during the Bronze Age and the Pre-Roman Iron Age. However, rye was not yet cultivated as a staple crop; instead, it primarily existed as a weed in fields of wheat (*Triticum* sp.) and barley (*Hordeum vulgare*), only becoming an established crop in its own right during the Roman Iron Age (Behre 1992). Pollen and archaeobotanical data from northern and northeastern Germany collectively show an increase in rye cultivation during the Roman Iron Age. However, throughout the LAMP there was a noticeable decrease, with an absence or very low presence of cereals, particularly rye, in both pollen and archaeobotanical records (Neef 2002; Behre 2008, p 253; Jahns et al 2013a; Jahns et al 2013b; Jahns 2018; Alinezhad et al 2024). Why did this decline occur? What triggered it? To answer these questions, we must consider the potential factors influencing cereal cultivation, including climate change, crop diseases, or their combined effects. This decline coincides with the climatic changes of the DACP and the LALIA. As shown in table 1, proxy-based climate data from various regions in Europe consistently indicate cold and wet conditions during the LAMP. These climatic shifts were characterized by lower temperatures in the growing-season and increased precipitation in northern and western Europe, which likely impacted agricultural productivity (Stamnes 2016; Büntgen et al 2016; Toohey et al 2018).

Beyond these climatic shifts, an enhanced vulnerability of cereals against diseases is for instance highlighted by Lee (2009) and Alm and Elvevåg (2013) and Bondeson and Bondesson (2014). These authors in two studies have documented that during 500-800 CE ergotism epidemics ravaged Europe due to the consumption of ergot-contaminated rye bread, resulting in a large number of deaths. Unlike prior studies from Scandinavia that primarily associate ergotism with the climatic anomalies of the 536 CE volcanic event (Alm and Elvevåg 2013; Bondeson and Bondesson 2014; Westling 2024), this study shifts the temporal focus from the 536 CE event to the broader DACP (350–700 CE) and the geographical focus from Scandinavia to northeastern Germany, as archaeobotanical evidence indicates an earlier presence of ergot in this region. Therefore, we use archaeobotanical data from domestic sites to assess cereal cultivation, particularly rye, before and during the DACP as well as the presence of ergot sclerotia in macrobotanical remains and its link to the broader historical processes including agricultural instability over multiple decades and subsequent human migrations of this period. We hypothesize that the cold, wet conditions of the DACP and the LALIA increased ergot (*C*. *purpurea*) infestation in rye, reducing crop yields and contributing to agricultural decline, which in turn drove socio-economic instability.

## 2. Methodology2.1 Palynological Data

The pollen data used in this study derive from lake Kleiner Tornowsee (KTO; 52.34°N, 14.05°E), located in Brandenburg, northeastern Germany (figure 1). This region, part of the North German Lowland, is characterized by a landscape shaped by Pleistocene glaciations and Holocene modifications, dominated by sandy plains with localized fertile areas (Stackebrandt and Franke 2015). Brandenburg has a subcontinental climate, with a mean annual temperature of 8.8°C and annual precipitation ranging between 500 and 560 mm. While the western part of Brandenburg has a more balanced maritime climate influenced by NAO variability, the eastern part is characterized by continental conditions with hot summers and cold winters (Unger and Lakes 2023).

**Figure 1.**
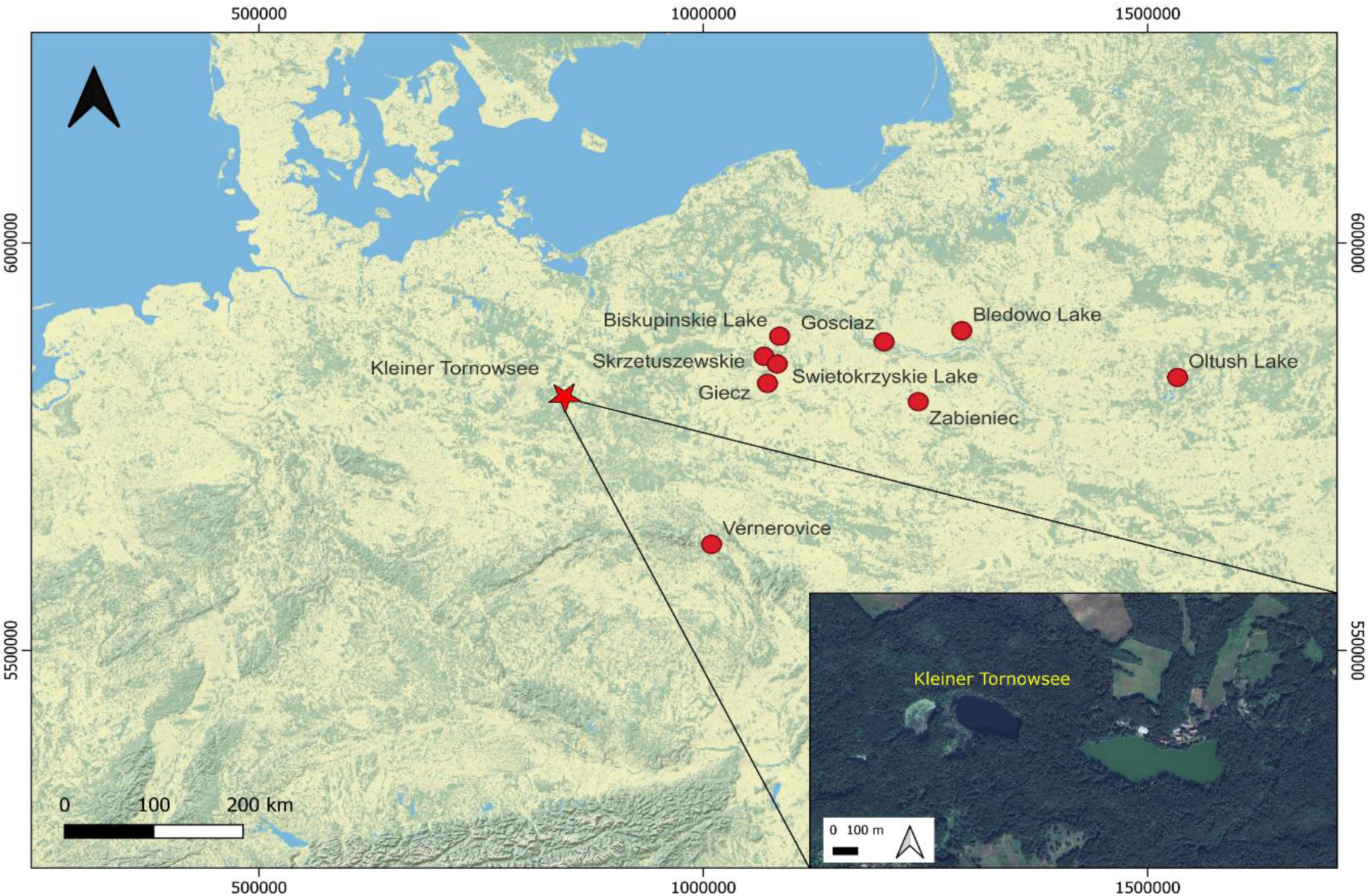
Overview map showing the location of lake Kleiner Tornowsee in the federal state of Brandenburg, Germany (KTO; marked with the red star) and the locations of other records (red dots) used for regional comparison

The high-resolution pollen record from lake KTO (figure 2), previously published by Alinezhad et al. (2024), is used here to examine new research questions. The chronology of KTO record is based on 31 radiocarbon dates (primarily terrestrial plant remains) and varve counts from the laminated section. The age-depth model was constructed using a Bayesian P_Sequence in OxCal 4.4 with the IntCal20 calibration curve. The modeled 2σ uncertainties range from –75 to +60 years, with a maximum of ±85 years at the top. This provides a robust chronological framework for the investigated period. The dates of the archaeobotanical and climatic datasets fall within this modeled uncertainty range, supporting their chronological consistency with the palynological record. Further methodological details on chronology can be found in Alinezhad et al (2024).

**Figure 2.**
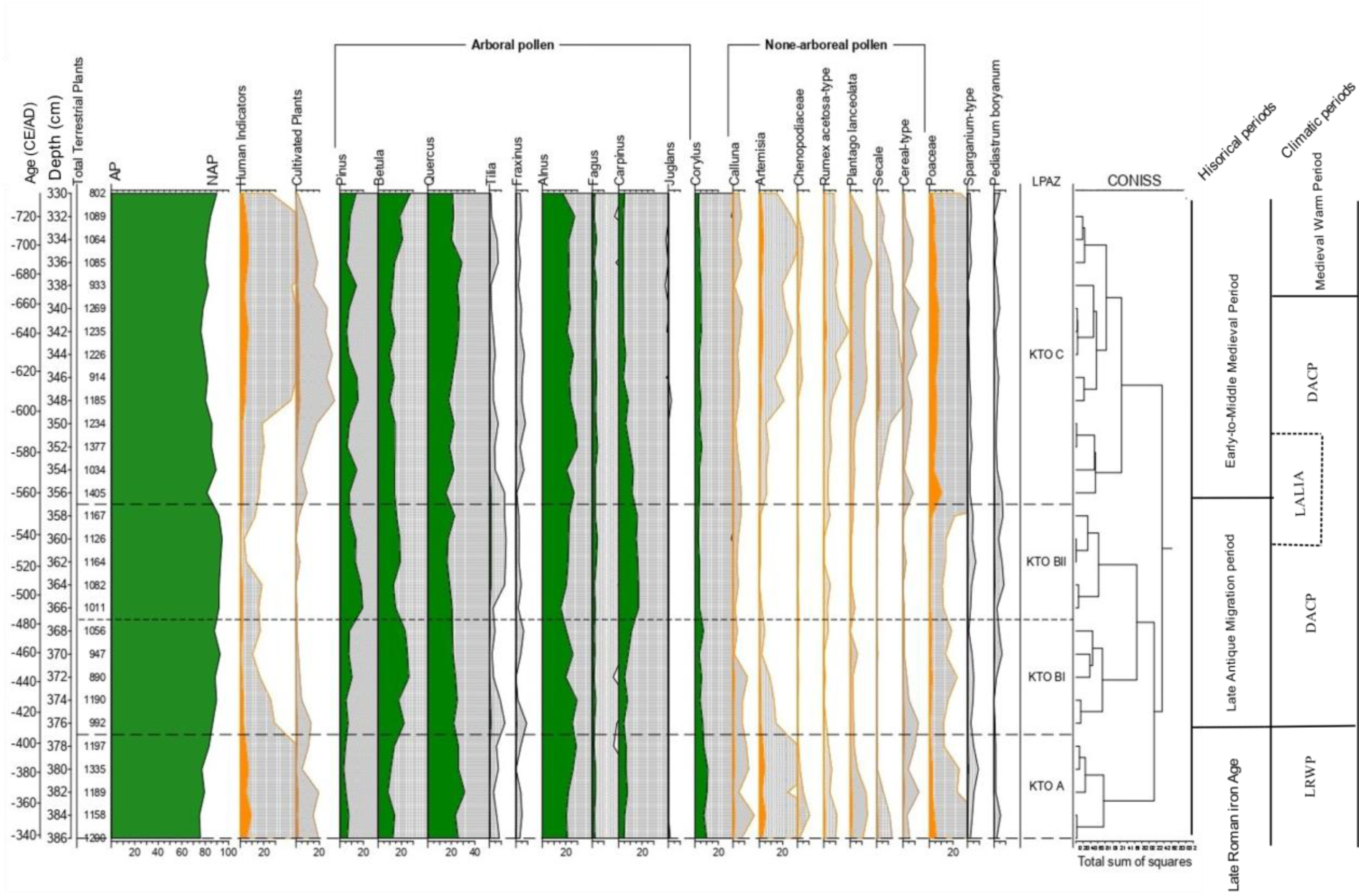
Simplified pollen percentage diagram of lake Kleiner Tornowsee (KTO) with 10-fold exaggeration indicated by grey silhouette; Historical periods (The Late Roman Iron Age, the Late Antique Migration Period, Early-to-Middle Medieval Period) and climatic periods (The Late Roman Warm Period (LRWP), The Dark Age Cold Period (DACP), The Late Antique Little Ice Age (LALIA), The Medieval Warm Period (MWP) are marked for chronological reference, showing vegetation changes and potential environmental drivers of land use changes (Alinezhad et al 2024)

The classification of cultivated plants and human indicators as anthropogenic indicators in Figure 2 follows Behre (1981). Cerealia and *Secale* (rye) are identified as cultivated plants, while *Plantago lanceolata* (ribwort plantain), *Rumex acetosa*-type (sorrel), Chenopodiaceae (goosefoot family), *Artemisia* (mugworts), *Centaurea cyanus* (cornflower), and *Calluna* (heather) are considered human indicators. These indicators were chosen for their connection with agriculture and settlement activities. The KTO pollen record, shows a pronounced decline in human activity, marked by reduced cereal cultivation (notably rye) and anthropogenic indicators, alongside forest regeneration during the LAMP. These trends align with broader archaeological evidence of settlement abandonment in northeastern Germany (Ejstrud et al 2008; cf. Volkmann 2014, p 137–138), but yet the drivers of this socio-environmental transformation remain unresolved. Using lake KTO’s well-established chronology and detailed record of vegetation and land-use changes, this study examines the interplay between climatic variability, cereal cultivation, and human migration during the LAMP. Specifically, we focus on the period ca. 340–700 CE to assess whether the cold and wet climate conditions of the DACP contributed to agricultural instability and, as a result, led to land-use abandonment.

To evaluate how cereal cultivation responded to climatic variability during the 4^th^-6^th^ centuries CE, we used cereal-type (*Hordeum* sp., *Triticum* sp., *Avena* sp.) and *Secale* pollen influx data (grains/cm²/year) from the lake KTO record (figure 3). Unlike relative pollen percentages, which can be affected by fluctuations in non-arable taxa (e.g., trees, shrubs, or weeds), influx values directly reflect variations in pollen accumulation rates (Davis 1963; Seppä and Hicks 2006). For instance, a decline in relative cereal pollen % could misleadingly suggest reduced cereal cultivation if tree pollen increases, even if actual cultivation remains stable. Further, the influx values reflect on-site cultivation rather than regional vegetation shifts (Andersen 1979; Faegri and Iversen 1989). Thus, influx data allows separating changes in crop productivity, enabling robust comparisons with annual temperature (TANN) and precipitation (PANN) reconstructions. This is important for detecting local agriculture activity as cereal-type pollen is poorly dispersed.

**Figure 3.**
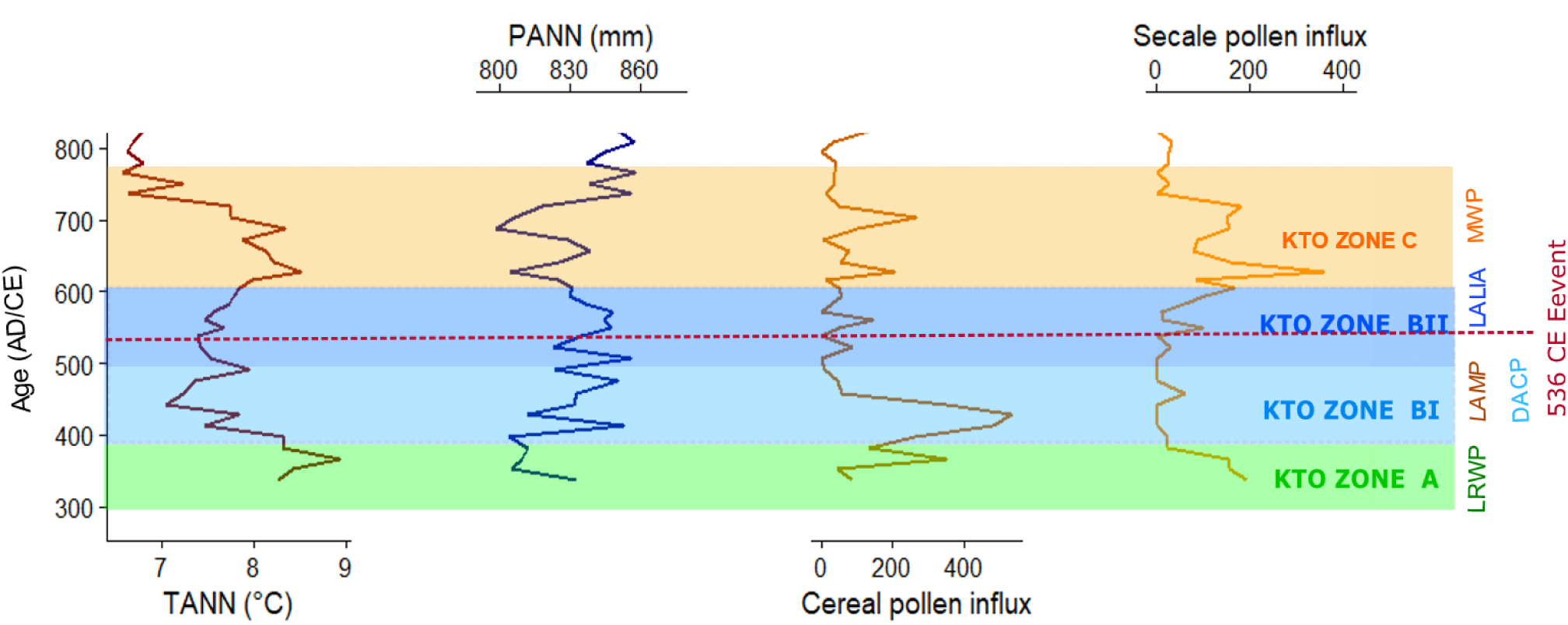
Pollen-based TANN and PANN estimates and cereal-type (*Hordeum* sp.*, Triticum* sp., *Avena* sp.) and *Secale* pollen influx derived from pollen data of lake Kleiner Tornowsee in northeastern Germany over the first millennium CE with focusing on 300-600 CE. The light green bar highlights the Late Roman Warm Period (LRWP; corresponding to KTO Zone A in the pollen diagram); the blue bars together highlight the cold and wet climate condition during the Dark Age Cold Period (DACP; corresponding to KTO Zones BI and BII in the pollen diagram) and the orange bar highlights the Medieval Warm Period (MWP; corresponding to KTO Zone C in the pollen diagram)

### 2.2 Climate Data

To assess the major climatic changes in NE Germany and NE Europe during the DACP, we used local and regional proxy-based climate reconstructions. In addition, climate model simulations from the Max Planck Institute Earth System Model (MPI-ESM), spanning 1 to 1850 CE (van Dijk et al 2022) were used to represent broader European climatic trends during the DACP. These simulations help place our local and regional climate reconstructions from NE Germany and NE Europe in a broader European context.

#### 2.2.1 Local and Regional Reconstructions

As the training set, a modern pollen dataset comprising 5,098 samples with a coverage of central and northern Europe (10° W - 44° E, 36° - 70° N) was used in this study. The modern pollen dataset was extracted from a previous study (Geng et al 2025). We integrated the modern pollen training dataset from Legacy Climate 1.0 (Herzschuh et al 2023b) and the EMPD2 (Davis et al 2020), which are both open-source. The training set from the modern pollen dataset within a 2000 km radius of the fossil sampling site was extracted. The fossil pollen dataset was extracted from lake KTO record (figure 3). The Weighted Averaging Partial Least Squares (WAPLS; ter Braak and Juggins 1993) method was performed as it is one of the most commonly used quantitative reconstruction methods for pollen-based climate reconstructions. WAPLS can handle non-linearity well and is more suitable than MAT to handle non-analogue situations. Therefore, we chose WAPLS as the statistical calibration approach to establish relationships between modern pollen data and climatic parameters for estimating paleoclimatic variables in R (R Core Team 2019) using the rioja package (Juggins 2022) and the reconstruction procedure was adapted from Geng et al (2024). After building the WAPLS reconstruction model based on the modern pollen training set, the fossil pollen record was then used to reconstruct climatic variables including annual temperature (TANN) and annual precipitation (PANN) (figure 3). The taxa names of the fossil pollen dataset were all harmonized in accordance with the modern pollen dataset. Only terrestrial pollen taxa were used, and rare taxa with values ≤ 0.1% were deleted, resulting in 34 taxa selected for analysis. A square root transformation of the pollen data was performed prior to all pollen-based analyses and reconstructions to stabilize variance and mitigate the impact of extreme values, such as abundant taxa or skewed distributions (Legendre and Legendre 2012).

The best component with the least error was identified using model statistics of WAPLS based on leave-one-out cross-validation and reconstruction uncertainties were indicated using Root Mean Squared Errors of Prediction (RMSEPs).

Moreover, the regional climate data (figure 4), which contains a subset (figure 1) extracted from a global compilation of temperature and precipitation data published by Kaufman et al (2020) and McKay et al (2024), were compared with the local PANN and TANN estimates based on pollen assemblages from lake Kleiner Tornowsee (figure 3). The coverage ranges from 51° to 54°N and from 10° to 20° E over the course of the last 12000 yrs. Temperature reconstructions are based on pollen assemblages from lakes and peats including one chironomid record using transfer techniques calibrated with modern training sets that provide quantitative estimates of past temperatures by correlating present-day distributions of proxy indicators with known climate variables. For precipitation reconstructions, data were drawn from McKay et al (2024), which provides a comprehensive analysis of hydroclimate variability during the Holocene. The study integrates 896 globally distributed datasets and employs a wide range of proxy records, including isotopic and sedimentary indicators, providing strong spatial and temporal resolution.

**Figure 4.**
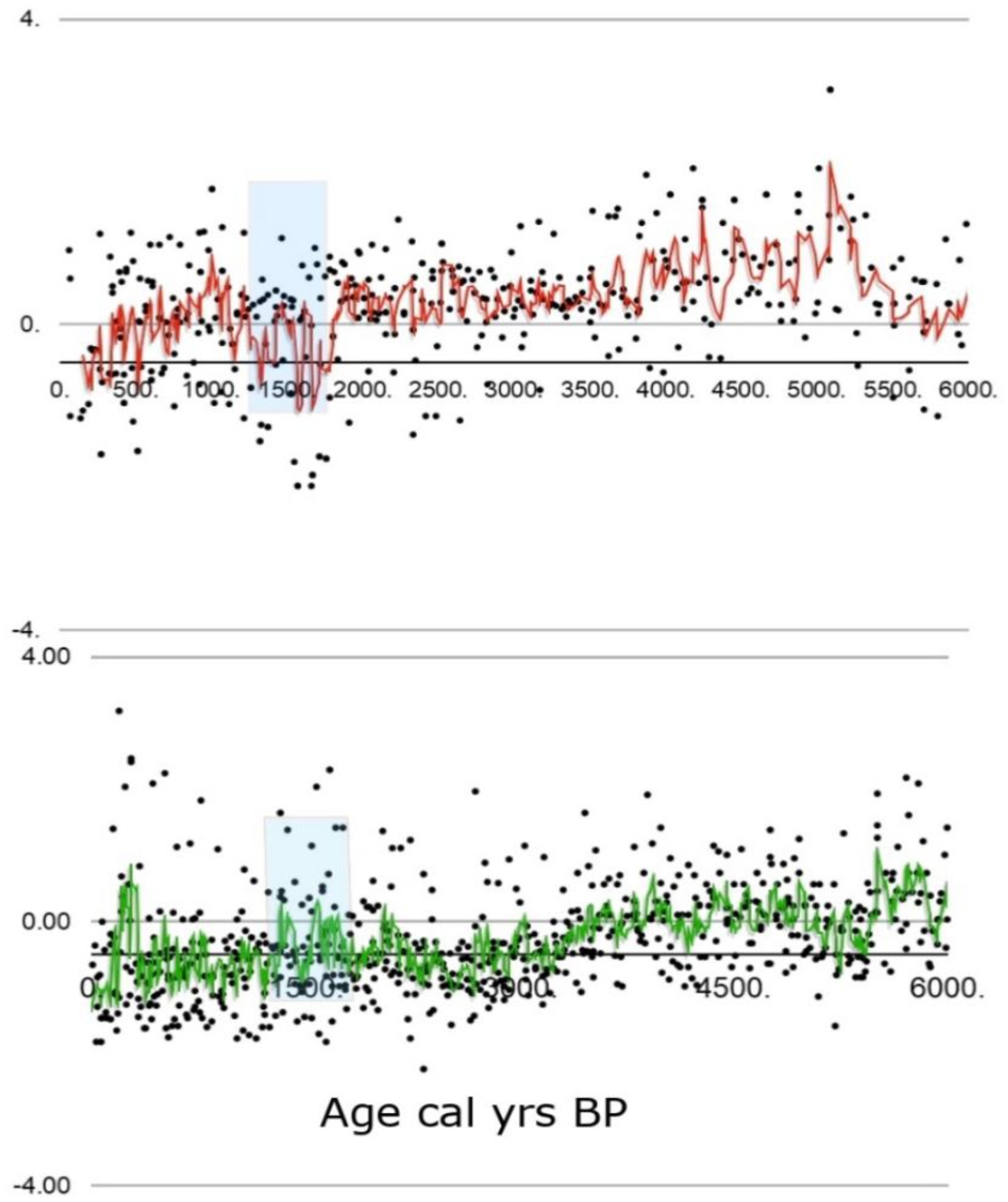
Reconstruction of the summer temperature (the upper plot) and the annual precipitation (the lower plot) for our study region. For summer temperature (50–55°N, 15–20°E, n=8), black dots represent the stacked z-score, and the red curve shows the 5-period moving average of the z-score. For annual precipitation (50–55°N, 15–20°E, n=10), black dots represent the z-score, and the green curve shows the 5-period moving average of the annual precipitation stack z-score. The blue bars indicate climatic trends and variability in both temperature and precipitation during the Dark Age Cold Period (DACP)

#### 2.2.2 Climate Modeling

To place our local and regional climate reconstructions results from NE Germany and NE Europe within a wider European framework, we compare them with climate model simulations from the Past2k project (van Dijk et al 2022). These simulations cover the period 1–1850 CE, and for this study, the period 300 CE to 700 CE is extracted to match our pollen records. This comparison helps us check whether the climate changes detected in the local and regional climate reconstruction results align with broader trends across Europe. The past2k runs are part of the Paleoclimate Model Intercomparison Project (PMIP) phase 4 as a contribution to the Climate Model Intercomparison phase 6 (CMIP6, Kageyama et al 2017). The aim of this experiment is to extend transient model simulations to the entire Common Era (CE), and to encourage model-proxy data comparison (Martrat et al 2019). The two past2k simulations were carried out using the MPI-ESM version 1.2. The atmospheric component (ECHAM6) has horizontal resolution of 1.9×1.9 degrees, with 47 vertical levels (top at 0.01 hPa, Stevens et al 2013). The ocean component (MPIOM) has a horizontal resolution of 1.5 degrees and 40 vertical levels (Jungclaus et al 2013). The full carbon cycle is included, represented by the submodels JSBACH and HAMOCC. The simulations are fully coupled transient runs, taken from the end of a 1200-year’spin-down’ simulation with constant year 1 CE boundary conditions. For more information on the model version, see Mauritsen et al (2019).

Since it is computationally expensive to carry out extensive fully coupled model simulations, only two simulations were executed for the past2k experiment with the MPI-ESM. Therefore, some climate signals could be hard to detect within the internal variability of the background climate, especially when covering smaller areas. This is amplified by the grid resolution of the model, which ranges between approximately 200 km at the equator, to 22 km near Greenland (Mauritsen et al 2019). Besides the grid resolution and the small number of simulations, parameterization of small-scale processes, and uncertainties in external forcing can also affect the model results. However, these are all well-known model biases, and unfortunately it is not yet feasible to carry out global, fully coupled earth system model simulations on a very small grid resolution. The MPI-ESM1.2 has been shown to perform quite well in studies for the CE (Timmreck et al 2021; Ward et al 2021; Fang et al 2022; 2023). In addition, the MPI-ESM past2k simulations have been shown to agree well with tree-ring records for the CE (van Dijk et al 2022). Multi-model intercomparison studies reveal that the MPI-ESM shows good agreement with other CMIP models regarding surface climate response on global and hemispheric scale (Hermanson et al 2020; Zanchettin et al 2022; Sospedra-Alfonso et al 2024). We therefore believe the MPI-ESM past2k simulations are a good choice for the model comparison part of this study.

To improve reliability, we study model anomalies over Europe (-10°E–30°E, 35°N–70°N; figure 5) rather than smaller regions like northeastern Germany (Brandenburg), as model data for very local areas can be less consistent and reliable. The average of two Past2k simulations is used to reduce random variations, and anomalies are calculated relative to the 1–1850 CE reference period, with significant changes identified using a 2σ threshold based on the full simulation period.

**Figure 5.**
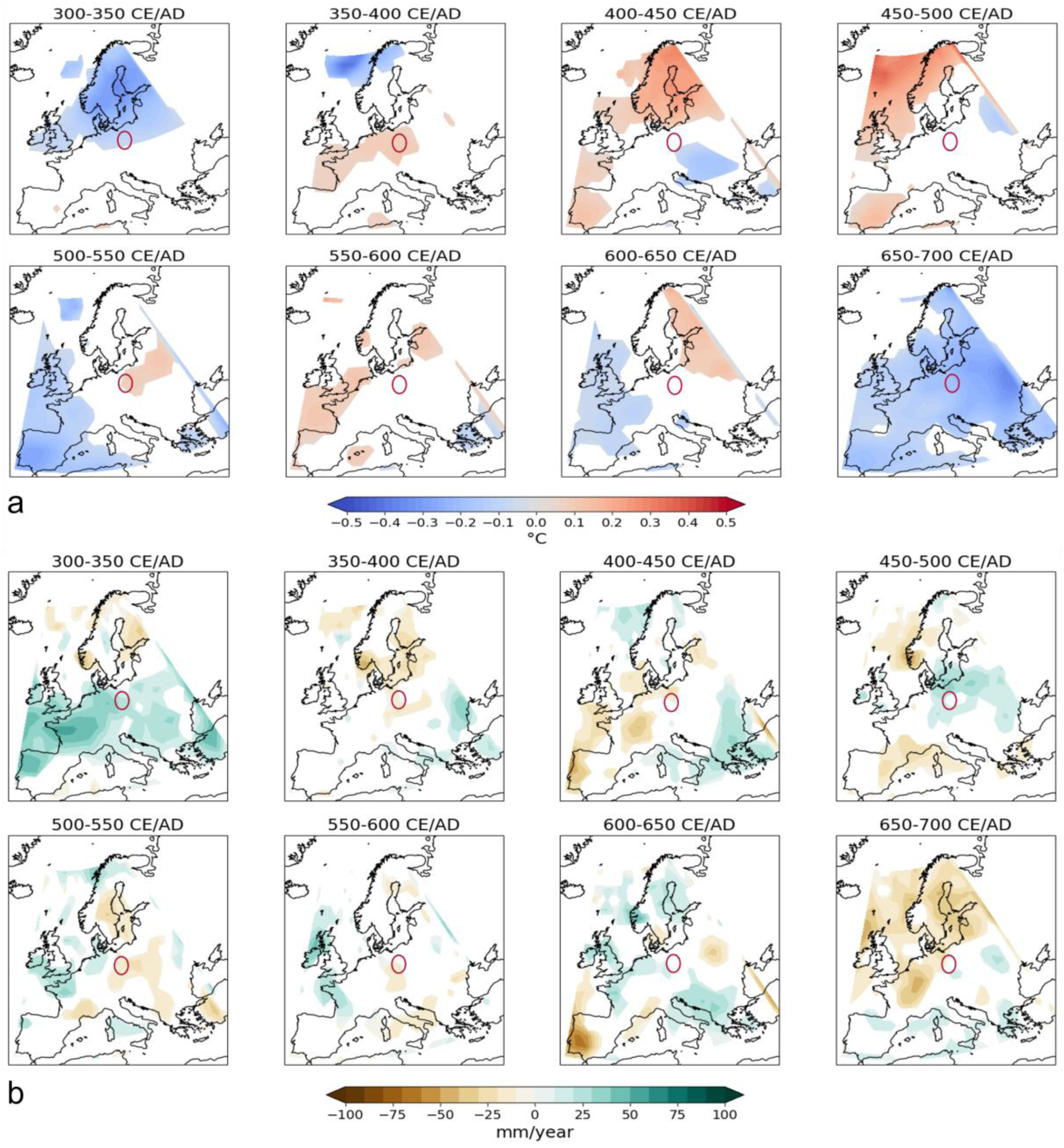
2m air temperature (a) and precipitation (b) anomalies for each 50-year mean during the DACP for Europe. Only values exceeding the 2σ threshold are presented; The red circle marks the Brandenburg area in northeastern Germany

The larger the area the reconstruction or simulation represents, the clearer external forcers leave a signal in the climate variables. Potential spatial biases exist when comparing local fossil pollen data with regional and continental-scale climate model data. These biases arise because each dataset measures and represents different types of data. The small size of Lake KTO (ca. 4 ha) makes its record highly sensitive to localized climate variability, and nearby land-use practices can subtly bias the pollen signal, factors well recognized in pollen-based reconstruction studies (Chevalier et al 2020). While these proxies remain robust for identifying broader climatic patterns, interpreting them requires caution and careful consideration of site-specific and anthropogenic influences. On the other hand, the model has a course spatial resolution, which leads to a smoothing effect over the grid space. In addition, some small-scale processes, such as cloud feedback, are parameterized rather than explicitly simulated. Combining datasets of different spatial scales is therefore challenging but offers valuable potential to disentangle genuine climate signals from internal variability and methodological biases.

### 2.3 Archaeobotanical Data

To assess cereal cultivation trends and their relationship to climatic and socio-environmental changes during the LAMP, archaeobotanical data on cereals from 14 archaeological sites in Germany, particularly from northeastern Germany (Brandenburg) (figure 6), were selected and added by notification on finds of ergot sclerotia when indicated. These datasets enable us to identify changes in cereal cultivation and possible connections to climatic stress during the DACP and cereal vulnerability to diseases such as ergotism, providing insights into agricultural instability and socio-environmental changes during the LAMP. Ergotism is a form of poisoning caused by the consumption of ergot-contaminated grains, particularly rye. Ergot (*Claviceps purpurea)* is a fungus that produces toxic alkaloids responsible for ergotism (Silva et al. 2023). More than 600 monocotyledonous plants, including rye, barley, wheat, oat, and wild grasses, are susceptible to this fungus (Schiff 2006; Berraies et al 2024). The production of ergot alkaloids depends on multiple factors, including the fungal strain, host plant species, temperature, humidity, and nutrient availability. Climatic conditions play a particularly influential role, as *C. purpurea* thrives in wet soils and high rainfall environments, which promote alkaloid production (Silva et al 2023). Rye is especially susceptible to ergot infestation among many host plants due to its wind-pollinated nature. Therefore, it was especially vulnerable during the 5^th^ and 6^th^ centuries CE when cold and wet conditions effected the flowering. In contrast, self-pollinating cereals like wheat were less susceptible to ergot due to more efficient fertilization (van Wyk 2005; Alm and Elvevåg 2013; Miedaner and Geiger 2015; Berraies et al 2024; Westling 2024).

**Figure 6.**
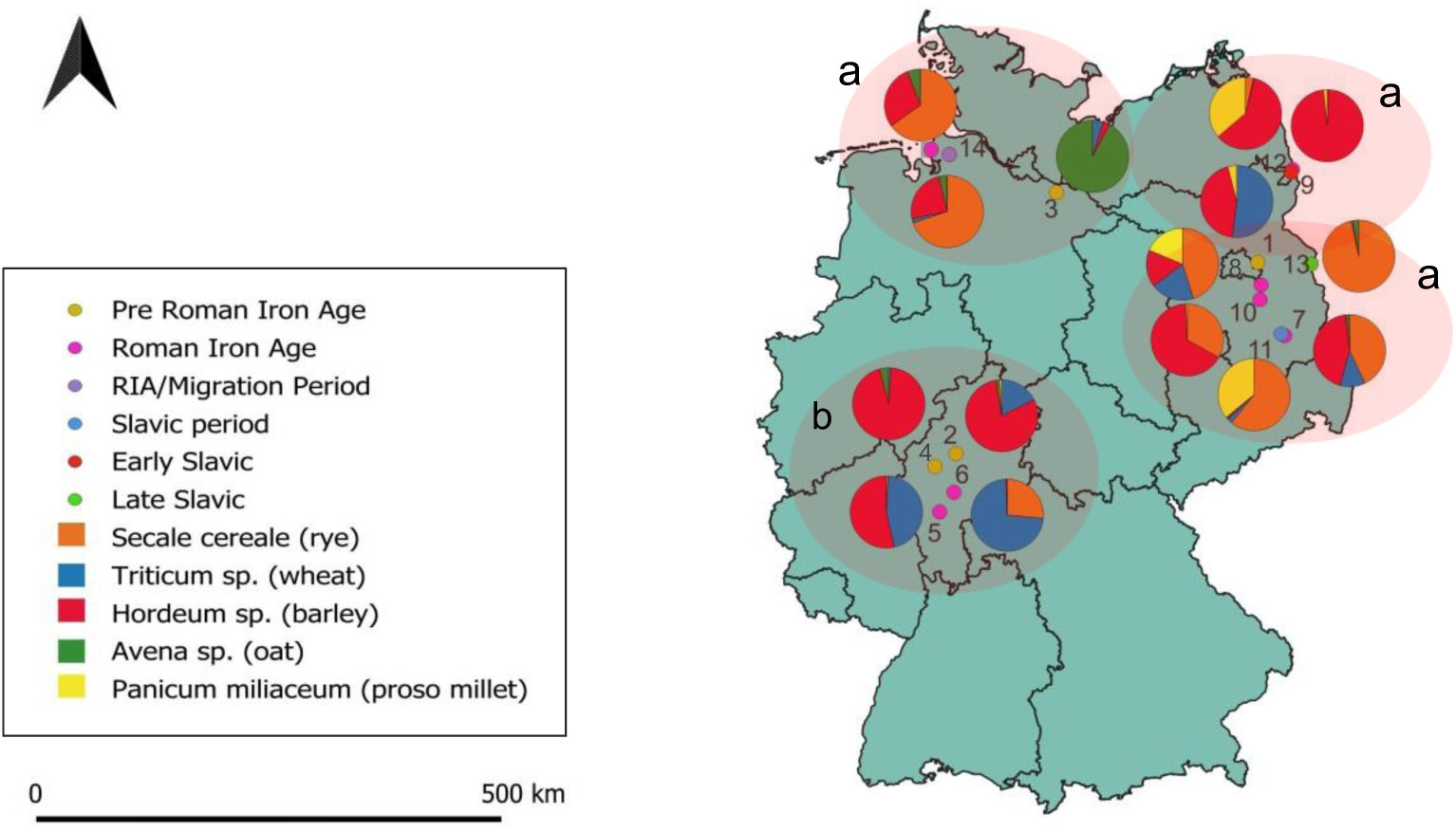
Archaeobotanical data on cereals from 14 archaeological sites across Germany, covering the Pre-Roman Iron Age (Neuenhagen, Mardorf, Rullstorf (1^st^ century BCE), Fellingshausen), the Roman Iron Age (Göritz, Klein Köris, Schwennenz, Kablow, Nieder-Eschbach, Echzell, Flögeln), the Migration Period (Flögeln) and Slavic period (Groß Lübbenau, Lebehn, Lebus). Pie charts show the proportions of major cereals: *Secale cereale* (rye), *Triticum* sp. (wheat), *Hordeum* sp. (barley), *Avena* sp. (oat), and *Panicum miliaceum* (proso millet). While pie chart sizes are uniform, the sites differ in total find quantities: 0–500 (Neuenhagen, Lebus); 500–5000 (Lebehn, Flögeln); >5000 (Echzell, Mardorf, Rullstorf, Fellingshausen, Nieder-Eschbach, Göritz, Klein Köris, Schwennenz, Kablow, Groß Lübbenau, Flögeln). Sites with sandy soils (circle a) include Neuenhagen (1), Rullstorf (3), Göritz (7), Klein Köris (8), Schwennenz (9), Kablow (10), Groß Lübbenau (11), Lebehn (12), and Flögeln (14). Sites with loess-derived soils (circle b) include Echzell (6), and those with fluvial gravels and aeolian sands (also circle b) are Mardorf (2), Fellingshausen (4), and Nieder-Eschbach (5). Lebus (13) is located on alluvial sediments. Soil information adapted from Feeser et al (2024, Abb. 1.2, p 5) after BGR (2016)

In regions with climatic and environmental conditions conducive to ergot outbreaks, such as colder, damp areas particularly in central, northern, northeastern, and eastern Europe—where rye cultivation was widespread (Prescott 1813; van Wyk 2005; Lee 2009; Bondeson and Bondesson 2014; Grikpėdis et al 2016), ergot infection was most common (Prescott 1813; Lee 2009). This is particularly significant given that rye thrives on dry, sandy soils and is sensitive to poor drainage soil (Westling 2024). One example is the evidence from archaeobotanical analyses of carbonized material from burnt houses in Sorte Muld on Bornholm and Vallhagar on Gotland that have demonstrated the occurrence of ergot in Scandinavia during the Migration Period (Helbæk 1957, p 272). The primary rye-growing region in Europe extended from the Low Countries through Germany to the loess areas of Central Europe, including present-day Czech Republic and Austria. Therefore, during the LAMP these parts of Europe were experiencing an agricultural instability and economic bust (Wickham 2009). The 14 archaeological sites from which the archaeobotanical data were used in this study fall within this broad region. These sites span four archaeological periods including the Pre-Roman Iron Age (PRIA), the Roman Iron Age (RIA), the Migration Period, and the Slavic (early and late Slavic phases) period. Four sites including Neuenhagen (Neef 2002), Mardorf (Wiethold et al 2008), Rullstorf (Behre 1990), and Fellingshausen (Kreuz 2004) are attributed to the Pre-Roman Iron Age. Seven sites date to the Roman Iron Age including Nieder-Eschbach, Echzell (Kreuz 2004), Göritz (Berg-Hobohm 2004), Klein Köris (Neef 2002), Schwennenz (Neef 2002), Kablow (Schiemann 1957). The site of Flögeln (Behre and Kucan 1994, p 54) spans both the Roman Iron Age and the Migration Period. The three remaining sites are associated with the Slavic period, comprising Groß Lübbenau (Medović 2004), Lebehn (early Slavic; Neef 2002), and Lebus (late Slavic; Neef 2002).

While the primary focus of this study is on cereal cultivation in northeastern Germany during the LAMP, data from earlier (Pre-Roman Iron Age and Roman Iron Age) and later (Slavic period) phases were included to provide a broader context for long-term agricultural trends. The archaeobotanical dataset used in this study was assembled according to criteria of data availability, geographic relevance, and accessibility of published sources. Due to limited access to archaeobotanical records from some nearby regions, the spatial scope was expanded to include additional areas of Germany, providing a broader context for interpreting cereal cultivation. While northeastern Germany, particularly Brandenburg, remained the primary focus in alignment with the regional scope of the pollen record, sites from central and northern Germany were also included. This broader perspective allows for the examination of whether cereal cultivation patterns observed in Brandenburg reflected regional or supra-regional trends, while also enabling comparisons across different ecological regions, such as the North German Plain with its sandy soils and the loess-dominated low mountain ranges of central Germany, as well as across different archaeological phases. The final set of 14 archaeological sites represents a curated dataset combining site-specific investigations and broader regional syntheses. The archaeobotanical methods reported in the source publications vary. More recent studies (e.g., Kreuz 2004; Wiethold et al. 2008; Medović 2004) employed systematic flotation techniques, whereas older studies, for instance, Schiemann (1957) relied largely on hand-collection and manual sieving of charred grains. Behre (1990) employed systematic water sieving followed by hand-sorting. Neef (2002) compiled results from multiple archaeological sites across eastern Germany, four out of which are considered in this study, although sampling methods are not always consistently documented. A lack of usage of sieves with small mesh size or hand-picking of solely large items as often encountered in early studies can be neglected for our analysis, since we focus on cereal and ergot finds, which belong to a size class above 1 mm. In addition, most of the assemblages derive from mass finds and storages, which enhances comparability.

For each site, we used the absolute find numbers of cereal grains as published in the original archaeobotanical studies: Neef (2002, table 25, p 322/323) for Neuenhagen, Klein Köris, Schwennenz, Lebehn, and Lebus; Wiethold et al. (2008, table 3, p 411; La2, La2/3) for Mardorf; Behre (1990, table 1, p 143; 1st century BCE) for Rullstorf; Kreuz (2004, table 12, p 137–144; AK1006 DÜN) for Fellingshausen/Dünsberg; Kreuz (2004, table 14, p 148–157; AK123 NES, RKZ2/3) for Nieder-Eschbach; Kreuz (2004, table 14, p 148–157; AK2569 EZ, RKZ 3) for Echzell; Berg-Hobohm (2004, table 50, p 142) for Göritz; Schiemann (1957, storage facilities D and E/F, pp 110 and 114) for Kablow; Medović (2004, table 2, p 192) for Groß Lübbenau; and Behre and Kucan (1994, table 8, p 54; 1st–3rd century CE) and (table 9, p 54; 4th–6th century CE) for Flögeln.

When calculating the sums, only whole cereal grains were included; double and triple identifications for cereal grains (e.g., emmer/spelt) were counted into the totals. Fragmented grains, chaff, and threshing remains were not considered. The compiled dataset with source references is provided in the supplementary material (Suppl. ## Archaeobotanical data).

Almost all sites show robust macrobotanical assemblages with 1,150 to 669,000 cereal grains recorded. Only two sites show lower find numbers, Lebus (n=348) and Neuenhagen (n=25), but still were included for the vicinity to the pollen site. To ensure comparability between datasets, we converted absolute counts into standardized ratios. The absolute find numbers for each cereal taxon were normalized by expressing them as a proportion of the total identified cereal macro-remains at each site. This method, as recommended by Popper (1988:214), helps account for differences in sample size and preservation by ensuring that each taxon’s representation is considered relative to the overall cereal assemblage. As a result, sites with lower preservation or smaller sample sizes are not disproportionately weighted compared to those with higher counts, enabling more meaningful cross-site comparisons.

Cereal species such as *Hordeum vulgare*, *Triticum aestivum, Avena sativa* are given under the genus-level classification *’Hordeum* sp.’, *’Triticum* sp.’, *’Avena* sp.’ and *Panicum miliaceum* and *Setaria italica* under’Millet’. This categorization is employed due to the constraints in pollen analysis, which typically allows for reliable identification only at the genus level. Therefore, to ensure consistency between the macroscopic botanical remains and the palynological data, all macro remains are grouped under the genus-level.

## 3. Results

### 3.1 Palynological Data

Pollen data from lake KTO in northeastern Germany (ca. 340–700 CE; Alinezhad et al 2024) provide high-resolution evidence of vegetation changes, cereal cultivation, and land use changes (figure 2). These data enable the investigation of human-environmental indicators during the Migration Period. The following section provides a comprehensive insight into vegetation patterns, human activities, and environmental changes based on palynological data (figure 2) from lake Kleiner Tornowsee in northeastern Germany during ca. 340-730 CE/AD (Alinezhad et al 2024). This pollen data is used in this study because they provide high-resolution evidence of shifts in cereal cultivation, land use, and anthropogenic activities, providing a suitable record for investigating human-environment interactions during the Migration Period.

Building on this dataset, our study aims to understand how changes in vegetation and human activities were influenced by environmental and climatic changes during the 4^th^-6^th^ centuries CE and whether these factors may have shaped human activities and contributed to human migrations in our study area. To address this question, we focus on anthropogenic indicators from the pollen record, such as cereal-type pollen and ruderal taxa, which reflect agricultural practices and settlement activities.

The pollen results show that between 340 and 400 CE, arboreal pollen (AP) dominates (70–80% of the total), while non-arboreal pollen (NAP) types such as Poaceae, *Artemisia*, *Plantago lanceolata*, and *Rumex acetosa*-type average 3–8%. Cereal-type pollen remains low (<2%), whereas *Secale* reaches ∼3% around 340 CE before declining. By 400 CE, *Secale* decreases further, while cereal-type pollen shows a slight increase (figure 2). From 400 to 480 CE, *Carpinus* expands, accompanied by a marked decline in anthropogenic indicators. Agricultural markers, particularly *Secale*, diminish, while pastoral indicators remain low, suggesting reduced human pressure on the landscape. Between 480 and 550 CE, woodland cover expands considerably (90–95%), with *Carpinus* reaching its maximum (15–20%). At the same time, anthropogenic indicators continue to decline, with some nearly disappearing. This points to a phase of widespread forest regeneration and minimal human activity. After 550 CE, arboreal pollen (AP) declines substantially (from 90–95% to 70–80%), with *Carpinus* falling sharply (15–20% to <5%) and *Betula* also decreasing. At the same time, non-arboreal pollen (NAP) increases, with *Secale*, *Artemisia*, *Plantago lanceolata*, *Rumex acetosa*-type, and Poaceae reaching higher values. This combination of reduced tree cover and expanding anthropogenic indicators reflects renewed settlement activity and agricultural expansion in the region.

### 3.2 Climate Data

The climate data for this study are reconstructed based on combined proxy-based climate reconstructions and climate model simulations, as described in section 2.2., which allow for a comparison between local, regional, and continental-scale climate trends during the DACP in Europe.

The local pollen-based climate reconstructions, based on the Lake Kleiner Tornowsee fossil pollen dataset (figure 3), reveal three distinct climatic phases between 300 and 700 CE. While the pollen-based climate reconstructions provide valuable insights into regional climatic trends, a potential minor distortion of the purely climatic signal due to local human impact cannot be entirely excluded. However, the general vegetation patterns are consistent with broader regional climatic trends, supporting the reliability of the reconstruction. During 300–390 CE, relatively low annual precipitation (PANN ∼810 mm) and moderate to high annual mean temperatures (TANN 8–9°C) coincide with high cereal-type and *Secale* pollen influx, suggesting stable agricultural conditions.

Between 390–580 CE, precipitation increases significantly (PANN peaks at 850 mm) while temperature declines by ∼2°C (TANN drops from 9°C to 7°C). This shift to cooler and wetter conditions coincides with an early decline in *Secale* pollen influx starting around 390 CE. Cereal-type pollen influx, on the other hand, shows a temporary increase, peaking around 430 CE, and then also declines. This pattern may suggest that the climatic and environmental changes initially impacted rye cultivation more strongly, leading to a short-term preference for other cereal crops. As unfavorable conditions persisted between 430 and 580 CE, overall cereal cultivation declined. Figure 4 illustrates the regional stacked climate data suggesting a drop in temperature (the upper plot) and a concurrent increase in precipitation (the lower plot) during the DACP, aligning with broader climatic trends of cool and wet conditions during this period in Europe. Although the datasets differ in temporal resolution—with temperature reflecting summer conditions and precipitation representing annual values— both datasets are relevant for assessing climatic conditions affecting vegetation and agriculture during the DACP.

To place these local and regional climatic patterns in a broader context, we examined climate model simulations from the MPI-ESM Past2k project (van Dijk et al 2022), which represent continental-scale climate trends.

The simulated 2m air temperature (figure 5a) and precipitation anomalies (figure 5b), presented as 50-year snapshots in figure 5, suggest that the DACP was not spatially or temporally homogeneous. In the first half of the 4^th^ century CE, Scandinavia experienced significant cooling, while mainland Europe showed anomalously wet conditions. From the second half of the 4^th^ century CE to the beginning of the 6^th^ century CE, only a few areas exhibited climate anomalies, and no clear pattern emerged. The first half of the 6^th^ century CE was anomalously cold in western Europe, with no significant hydroclimatic changes. By the second half of the 7^th^ century CE, cooling extended across all of Europe, with dry conditions prevailing in Northern Europe and Scandinavia.

The discrepancies between the model simulations and proxy reconstructions are due to a number of causes, which are challenging to disentangle. Part of the discrepancies stem from the differences in spatial resolution, where the model is smoothing the output over the entire grid cell, which covers a larger area. This means that the model cannot resolve small-scale processes properly. Other uncertainties related to the model stem from the background climate. Slight changes in the starting climate of the model simulation impact its evolution (the butterfly effect). A large ensemble could overcome these issues, but unfortunately, this is not feasible for such long simulations, as they are computationally expensive. On the proxy side, chronological uncertainties in the pollen age-depth model may distort the timing of reconstructed trends. In addition, because the lake covers a relatively small area, the record may reflect localized climate variability and may also contain anthropogenic signals that are not represented by the model.

### 3.3 Archaeobotanical Data

To provide a visualization of the distribution and proportions of various cereal data from different archaeological sites in Germany during the Pre-Roman Iron Age (ca. 800-1 BCE/BC), Roman Iron Age (ca. 1-500 CE), and the Early and the Late Slavic periods (ca. 600-1200 CE), the available archaeobotanical data on the cereals from 14 terrestrial sites with charred plant preservation (data compilation in the supplementary material) are presented in pie charts displayed on the map (figure 6).

The map (figure 6) illustrates the spatial and temporal variations in cereal assemblages, providing insight into cereal cultivation practices across different regions of Germany during these periods. The dataset is spatially and temporally uneven, with limited coverage for some regions and for the Migration Period. Additionally, when interpreting the data, it is crucial to keep in mind its limitations: the charred material provides us with a snapshot of what crops where available at the respective site in a burning event rather than being fully representative for the sites plant economy. Nevertheless, this compilation allows addressing possible shifts in cereal cultivation and the presence of ergot prior to and after the DACP, improving the classification related to the examination of the research hypothesis that the cool and wet conditions of the DACP may have fostered ergot growth, likely reducing agricultural productivity. Therefore, to better understand this relationship, the archaeobotanical data were examined for changes in cereal crop composition, especially rye, alongside the frequency of ergot sclerotia before (Pre-Roman and Roman Iron Age), during (Migration Period ca. 400–600 CE), and after (Slavic period) the DACP. The survival of ergot sclerotia in archaeobotanical remains depends on specific preservation conditions. The delicate, alkaloid-rich sclerotia (dormant structures) break down rapidly unless preserved by charring or in anaerobic environments, so they are typically found at sites with these preservation conditions (Heiss et al 2017). Additionally, ancient crop processing methods (e.g. sieving, winnowing, flotation) could likely remove the sclerotia. Furthermore, modern macro-remain processing techniques may unintentionally overlook ergot remains, potentially explaining gaps in our understanding of ergot’s impact on rye cultivation in the past (Barclay and Fairweather 1984; Jones 2012). Thus, while finding ergot sclerotia provides valuable insights into historical crop diseases, their absence does not necessarily indicate a lack of infestation.

Seven sites from the RIA and MP dating to the 3^rd^ to 5^th^ centuries CE show evidence for ergot sclerotia. The increased presence of ergot in these archaeobotanical assemblages coincides with climatic stress during the DACP and a greater reliance on rye cultivation in northeastern Germany. This pattern supports the broader hypothesis that the DACP contributed to agricultural instability, potentially leading to population decline and changes in land use. Although ergot finds remain limited and cannot be used to quantify the scale of infestation or its direct impact on demographic shifts, their temporal and ecological alignment with the DACP conditions strengthens the argument that ergot proliferation was a contributing factor in climate-induced agricultural stress during the Migration Period.

### Pre-Roman Iron Age (Sites 1–4)

From the pre-Roman Iron Age, there are only a few macrobotanical studies available from the old moraine (the Saalian glacial landscapes, which are strongly weathered, leached, and nutrient-poor) in the North of Germany (old moraine landscapes). Rullstorf (3; Behre 1990), a site with sound archaeobotanical data derived from storage pits dating to the 1^st^ century BCE with a total of 6,649 cereal finds, presents oat dominance (92%), while barley and wheat are minor (3% and 5% respectively), and rye is absent. In Neuenhagen (1; Neef 2002), both wheat and barley are present, while rye is absent. However, the representativity of findings from this site is very limited due to the very small number of cereal macro-remains (n = 25). The source publication does not provide detailed information on taphonomy or excavation conditions, but the low counts may result from poor preservation, limited sampling area, or excavation strategy.

On the loess-dominated soils in the loess plains of central Germany, barley and wheat are the main crops during the Pre-Roman Iron Age. In the basaltic and metamorphic soil region at Mardorf (2; Wiethold et al 2008), with 37,833 cereal remains recovered from waste pits and postholes, barley is dominant (80%), followed by wheat (18%) and minor oat (2%). Similarly, in Fellingshausen (4; Kreuz 2004), with a total of 13,555 cereal remains from a pit, barley is the dominant crop (95%), with oat playing a minor role (4%).

Overall, wheat, barley, and oat were cultivated across all regions. However, the dominant crops differed between regions, with barley and wheat being more prominent in the loess areas, and barley prevailing in the basaltic and metamorphic soils.

In addition to these crop patterns, occasional traces of ergot are also present. Ergot finds have been documented in Rullstorf (Early Pre-Roman Iron Age; 6^th^-2^nd^ centuries BCE) and Mardorf (Pre-Roman Iron Age).

### Roman Iron Age (Sites 5–10, 14)

In northern Germany during the Roman Iron Age, Flögeln (14; Behre and Kucan 1994, p 54), with 1,155 cereal remains, shows notable cultivation of rye (70%) alongside secondary barley (24%) and minor oat (4%) while wheat and common millet are nearly absent. In northern and northeastern Germany, where rye was largely absent during the Pre-Roman Iron Age, a significant shift occurs with the emergence of rye in large quantities during the Roman Iron Age. On the sandy soils of the young moraine in Göritz (7; Berg-Hobohm 2004, p 142), with 81,704 cereal grains from pits, and Kablow (10; Schiemann 1957, 110 and 114), with 29,440 cereal grains from storage pits, rye (43% and 45% respectively) is a main cereal with barley (44%) as second main cereal in Göritz and wheat (20%) in Kablow. In Klein Köris (8; Neef 2002), with 22,067 cereal grains, barley is dominant (66%), with rye as the secondary cereal (33%) and minor amounts of millet (1%) and wheat and oat are almost absent.

On the sandy soils of the old moraine in Schwennenz (9; Neef 2002), with 299,508 cereal remains recovered from pits and postholes, barley is the dominant crop (98%), rye, oat, and wheat are also present but in minor quantities (less than 1%), and millet appears as well in low quantities (2%).

In Nieder-Eschbach (5; Kreuz 2004), in central Germany on soils with fluvial gravels and aeolian sands, 103,364 cereal grains were recovered from storage contexts. This site is dominated by barley (53%), with wheat (46%) as a secondary cereal. Rye is present only in minor amounts (2%), oat is absent, and common millet occurs in very small quantities (less than 1%). On the loess-derived soils in Echzell (6; Kreuz 2004), with 6,376 cereal remains from storage, wheat is the dominant crop (73%) and rye (26%) is secondary cereal. Barley is present in minor amounts (1%) and oat and millet are absent.

Despite regional variability, a broader pattern emerges in which rye appears ubiquitously but with varying dominance. In Göritz and Kablow, rye is a main cereal, while in others, like Schwennenz, Klein Köris, and Echzell, it is secondary to barley or wheat. Furthermore, ergot sclerotia are found in the moraine areas, such as Göritz, Schwennenz and Kablow, primarily at sites where rye is a main cereal.

### RIA/Migration Period (Site 14)

Finds from the Migration Period from central and northeastern Germany are almost lacking. However, the Migration period is recorded at the multi-period site of Flögeln, on the sandy soils of the old moraine in northern Germany. At Flögeln (Site 14; Behre and Kucan 1994, p 54), 35,902 cereal grains were recovered from storage pits, postholes, a kiln, and cultural layers. Rye dominates the assemblage (65%), followed by barley as a secondary cereal (29%). Three finds of ergot were also recoded at the site, dating to the Migration Period.

### Slavic Period (Sites 11–13)

On sandy soils of the young moraine in Groß Lübbenau, a large assemblage of charred plant remains from a cultural layer left after a massive fire event in the Slavic fortified settlement with 669,021 cereal grains was recovered (11; Medović 2004, p 192), and on sandy soils of the young moraine in Lebus (13; Neef 2002). Rye dominates the cereal grains (61% and 97% respectively), followed by secondary proso millet (36%) and minor quantities of wheat (2%) in Groß Lübbenau and oat (2%) in Lebus. However, it should be noted, the data from Lebus are quite scarce, consisting of only 348 cereal grains and the archaeological context has not been detailed in the publication. In contrast, on the young moraine at Lebehn (12; Neef 2002), a settlement site with archaeobotanical samples from pits, with a total of 2,993 cereal grains, rye is present in minor quantities (4%) while barley dominates (60%) and proso millet (36%) is secondary. Despite such local variations, the broader trajectory points toward an increasing reliance on rye cultivation. In eastern Germany, the Slavic period marks the full ascendancy of rye as the dominant cereal crop from 1000 CE onwards. Findings of ergot are recorded solely in the site Groß Lübbenau (Medović 2004, p 192).

The current state of the archaeobotanical data remains limited. The very small number of sites dating to the LAMP, represented only by Flögeln for the Migration Period, hinders direct inferences about crop cultivation and environmental constraints. The sparse data may suggest reduced agricultural activity, though further evidence is needed to confirm this. Despite this fact, the data shows the growing importance of rye from the RIA onwards. It is probable that its relevance was growing throughout the LAMP, although its full dominance as the main crop does not occur prior to 1000 CE (Behre 1992; Behre 2008). Regional patterns suggest that rye was dominantly cultivated on sandy soils, especially in the young moraine landscapes of northeastern Germany (e.g., Kablow) and on the sandy soils of the old moraine in northern Germany (e.g., Flögeln), while barley tended to dominate in areas with loess, fluvial, or wind-blown deposits, such as Fellingshausen or Nieder-Eschbach. The regular notice of ergot sclerotia at rye dominant sites of moraine regions further makes it probable that ergot infestation of rye had an effect on cereal yields during the LAMP. The increasing presence of rye and its co-occurrence with ergot sclerotia, particularly during the cold and wet climatic conditions of the DACP, support the hypothesis that these environmental changes created favorable conditions for ergot outbreaks in northeastern Germany. We therefore interpret this link as a plausible hypothesis supported by broader discussions of ergot’s historical role in agricultural instability (Alm and Elvevåg 2013; Bondeson and Bondesson 2014), while emphasizing that further research is required to assess its scale and impact.

## 4. Discussion

### 4.1 Dark Age Cold Period Climate Change, Environmental Changes, and Agricultural Instability during 4^th^-7^th^ Centuries CE

In the Late Holocene, Europe experienced several centennial-scale climate fluctuations, such as the Roman Warm Period (RWP), the Dark Ages Cold Period (DACP), and the Medieval Warm Period (MWP), all of which significantly influenced past societies and environmental conditions (Waltgenbach et al 2021). The RWP (300 BCE to CE 350) in Europe was followed by the DACP (350 to 700 CE). Proxy data from the DACP collectively indicate prevailing cold and wet conditions in Europe (table 1), likely with similar temperature anomalies (Wanner 2008; Helama et al 2017; Helama 2021; Waltgenbach et al 2021; McKay et al 2024).

The syntheses of the pollen-based climate reconstructions in Europe during the Holocene indicate that both the annual and summer temperature anomalies show a distinct trough around 450 CE (1500 cal yrs BP) (Geng et al 2025). This observation at the pan-European scale also coincides with the reconstruction of the summer temperature for our study region. Figure 4 reveals a sequence of reconstructed climatic patterns in 50-year intervals for the broader study region, specifically summer temperatures and annual precipitation. A clear cooling anomaly is apparent around 400 CE (1550 cal yrs BP), corresponding to ca. 390–580 CE, marked by lower summer temperatures compared to both preceding and subsequent periods. Simultaneously, annual precipitation increases during the same interval, indicating wetter climatic conditions. Together, these continental and regional observations confirm that the broader regional climatic setting during the DACP was characterized by cool and wet conditions, which align well with our pollen-based local reconstructions from lake Kleiner Tornowsee (figure 3).

Comparative data from regional pollen records, including Sacrower See (Alinezhad et al 2024), Wittwesee (Jahns unpubl.), and Rudower See (Jahns et al 2013a; Jahns et al 2013b; Jahns et al 2018), consistently demonstrate similar vegetation responses—marked by widespread woodland expansion, notably hornbeam (*Carpinus*), and reduced human agricultural activities. This reduction in human activities is mirrored in the archaeological record, which shows that from the late 4^th^ century onward, northeastern Germany experienced phases of settlement abandonment. These demographic shifts have been linked to the westward and southward migration of Germanic groups (Volkmann 2014, p 137– 138). This regional consistency suggests that the climatic changes during the DACP and the LALIA broadly impacted agricultural productivity and human settlement patterns across northeastern Germany and neighboring regions in Poland and the Netherlands (Rösch 1992, 1993; Ralska-Szpiganowicz et al 2004; Dreßler et al 2006; Czerwiński et al 2022; Pierik 2021; Lukanina et al 2023; Gouw-Bouman 2023).

The cooler, wetter DACP climate likely shortened growing seasons and made cereal cultivation unstable. In addition to political upheavals, the decline in cultivation productivity and resulting agricultural instability likely pushed local populations to migrate or abandon their farmlands. Helama et al (2017) highlight that colder temperature and increased moisture, especially during the spring and summer seasons, would have reduced the growing periods and yields of crops such as cereals, which are highly sensitive to seasonal changes. Seasonality played a crucial role, with spring and summer conditions becoming wetter, which likely disrupted planting and harvest cycles, further aggravating agricultural instability (Helama et al 2009; Swierczynski et al 2012; Grauel et al 2013). Moreover, Helama et al (2017) suggest that the cold conditions of the DACP were driven by a combination of factors, including a negative phase of the North Atlantic Oscillation (NAO), reduced solar activity, and ice-rafting events in the North Atlantic, all of which contributed to prolonged cold winters and cooler summers. These climatic changes likely made cereal farming unsustainable in many regions of Europe including northeastern Germany, leading to the observed decline in agricultural activity and the corresponding expansion of woodlands, particularly hornbeam.

While our regional and site-specific evidence indicates a consistent pattern of cooler and wetter conditions during the DACP, Helama et al. (2017) emphasize that although many records documents cooling between ca. 400 and 765 CE, the climatic expression of the DACP was not uniform. Northern and central Europe show cold/wet signals, whereas Mediterranean and Tibetan Plateau records indicate coincident droughts. Similarly, Grauel et al (2013) reconstruct wetter conditions in the Gulf of Taranto (central Mediterranean) between 500 and 750 CE, but within a broader context of gradual drying across the wider Mediterranean basin. Moreover, the DACP itself remains difficult to define precisely, given the lack of consensus on its trigger, onset, and duration (Esper et al 2014; Luterbacher et al 2016). Although increased climate variability is recognized in the 4th and 5th centuries CE, abrupt changes and clear forcing mechanisms have not yet been identified (Helama et al 2017). Climate simulations further support the view that the DACP was not a uniform climatic regime but rather a temporally and spatially unstable interval. This variability highlights that, although our study region shows a coherent cool and wet pattern, broader European and extra-European evidence underscores spatially varied expression of the DACP. At the same time, our KTO record, because of its small size (ca. 4 ha), primarily reflects local vegetation dynamics. However, comparison with nearby regional pollen records from northeastern Germany shows parallel vegetation and land-use patterns and thus supports the interpretation that the KTO record also captures broader sub-regional trends. Using pollen data for local climate reconstructions is a reliable and well-established method, though the record may also contain subtle signals of human land use. In addition, chronological uncertainties in the age–depth model (with 2σ ranges of up to ±85 years) could affect the chronological alignment of reconstructed climate phases, which we considered when we compare local, regional, and continental-scale datasets.

### 4.2 Vegetation and Demographic Changes, Human Activity Decline during 4^th^-7^th^ Centuries CE

The pollen results from ca. 340 CE (2σ range of 260-390) to 400 CE (2σ range of 330-460) provide insights into the vegetation composition and the anthropogenic activities in the northeastern Germany during the Late Roman Iron Age, a period when it was inhabited by Germanic groups. The results suggest the dominance of woodland, with evidence of cereal cultivation, particularly rye, and indicators of pasture and fallow-land such as *Plantago lanceolata*-type and *Rumex acetosa*-type (Behre 1981), signifying agro-pastoral activity in the region during this time. This evidence points to a relatively dense settlement by the Germanic groups who were skilled at utilizing the available resources to establish and sustain an agro-pastoral economy (Hamerow et al 2023). They had livestock grazing in the floodplains or in relatively open parts of the landscape, and they practiced agriculture on arable fields.

After the Late Roman Iron Age (ca. 340-400 CE), the pollen results from ca. 400 CE (2σ range of 330-460) to 480 (2σ range of 410-540) show a notable decline in anthropogenic indicators particularly agricultural indicators such as *Secale*, while there is still a low presence of pastoral indicators visible. These changes are followed by a gradual increase in woodland, especially in hornbeam. The decrease in anthropogenic indicators and the beginning of woodland expansion in ca. 400 CE (2σ range of 330-460) to 480 CE (2σ range of 410-540), coincides with the first phase of Germanic migration from the territory of the modern state of Brandenburg in western and southern directions. Archaeologically, a gradual decline of settlements can be observed from at least the late 4^th^ century CE onwards, which is usually associated with migration towards provinces of the Roman Empire, particularly to Great Britain (cf. Volkmann 2014, p 137-138).

After the onset of human activity decline and woodland expansion during the early Migration Period (ca. 400-480 CE), AP increased, reaching its highest levels (95-98%). *Carpinus* reached its maximum distribution and anthropogenic indicators especially pastoral and fallow-land indicators such as *Plantago lanceolata*-type and *Rumex acetosa*-type declined to low values between ca. 480 CE (2σ range of 410-540) to 550 CE (2σ range of 480-610). The marked expansion of *Carpinus* between ca. 480 and 550 CE, suggests that woodland regeneration during the Migration Period was not simply a passive consequence of human abandonment (Rösch 1992; 1993; Ralska-Jasiewiczowa et al 2004; Dreßler et al 2006; Pierik 2021; Czerwiński et al 2022; Lukanina et al 2023). Instead, ecological processes played a role in shaping forest composition. Two factors in particular may explain the success of *Carpinus*. Firstly, its capacity to resprout vigorously from stumps after coppicing which allows it to recover quickly on previously managed land, and secondly the prevalence of cold winters during this period (Helama et al 2017), which may have given hornbeam a competitive advantage over more climate-sensitive taxa such as *Fagus*. This interpretation highlights that woodland expansion resulted from the interaction of both reduced land use and ecological succession, with *Carpinus* emerging as a key beneficiary of these dynamics.

The sharper decline of anthropogenic indicators, compared to the former phase of abandonment, ca. 400 CE (2σ range of 330-460) to 480 CE (2σ range of 410-540), highlights the greater intensity of the Germanic abandonment in this phase. It is during the latter half of the 5^th^ century CE, that a significant number of people migrated from the area of northern Germany, resulting in a significant rise in Anglo-Saxon settlers across eastern and southern England (Clarke 1985, p 40; Ejstrud et al 2008).

After the period of settlement abandonment from around 400 CE (2σ range from 330-460) to approximately 550 CE (2σ range of 480-610), significant changes in both human activity and vegetation composition are evident in the study area between 550 CE (2σ range of 480-610) and 730 CE (2σ range of 650-780). During this time, the decline in arboreal pollen, especially *Carpinus* and *Betula*, is accompanied by an increase in *Quercus*. This shift coincides with a notable rise in cultivated crops such as cereal-type and *Secale*, alongside secondary indicators of human activity like *Plantago lanceolata*-type, *Rumex acetosa*-type, *Artemisia*, and Poaceae, which suggest the presence of both, woodland pasture as well as arable farming around the lakes. These agricultural and pastoral practices played a key role in shaping the region’s landscape.

To understand the contributors underlying the aforementioned vegetation and demographic changes during the LAMP, several factors need to be considered. Archaeologically, one contributing reason may have been the Roman withdrawal in the early 5^th^ century CE, leaving England vulnerable to attacks from groups like the Picts and Scots. In the mid-5^th^ century CE, Britons sought the assistance of Germanic mercenaries to defend their land, allowing them to settle in East Anglia, where better living conditions were available (pull factors) (cf. Volkmann 2014, p 137-138). Another factor could have been climatic and environmental changes, along with agricultural instability, that caused economic hardships (push factors) (Büntgen et al 2016; Schneeweiß 2023). Climate data indicate that the onset of cooler and wetter conditions in the 5th century CE, associated with the DACP and later intensified during the LALIA, shortened the growing season and increased hydroclimatic variability (Büntgen et al 2016; Toohey et al 2018; Helama 2021). These shifts reduced harvests, undermined cereal cultivation, and created agricultural instability that threatened food security, which could have forced settlements to seek more secure land and resources. Climatic stress and its agricultural consequences therefore acted as a trigger which, in interaction with political instability and social dynamics, contributed to the large-scale migrations of the 5th century CE (Schneeweiß 2023). Rather than being the only driver, these environmental pressures amplified existing vulnerabilities and contributed to widespread human mobility during the Migration Period.

These findings from Brandenburg fit into a broader European pattern of the Migration Period disruptions. In Scandinavia, the 5th–6th centuries saw widespread depopulation and forest regrowth, often linked to climatic downturns and the 536 CE event (Gräslund & Price 2012; Alm & Elvevåg 2013). Across central Europe, palaeoecological studies show parallel woodland expansion and declining anthropogenic indicators during this time (Dreßler et al 2006; Czerwiński et al 2022). Similar, in western Europe, the migration of Germanic groups into Roman provinces coincided with opportunities for more stable subsistence economies (Volkmann 2014; Clarke 1985). Germanic groups were recruited as mercenaries by the Britons and were allowed to establish themselves in regions where they could access fertile land and better living conditions that offered advantages compared to their areas of origin (Volkmann 2014). Therefore, to avoid oversimplification, it is important, alongside the social and political reasons, to investigate and discuss these environmental and climatic factors as potential contributors, which will be further explored in the following paragraphs.

### 4.3 Rye Cultivation and Ergot Infestation: A Consequence of Climatic Stress?

Due to limited data, the following interpretations should be considered provisional. Nevertheless, the review of archaeobotanical evidence indicates that cereal cultivation was well-established during the Pre-Roman Iron Age in Germany; however, rye is notably absent in northern and north-eastern sites (figure 6). According to Behre (1992), the onset of rye cultivation can be traced back to the Pre-Roman Iron Age in Central Europe and it became one of the most important crops alongside barley on the poor moraine soils (these soils were formed by the deposition of sandy sediments by glaciers during the last Ice Age, known as the Pleistocene era) during the Roman Iron Age. A further massive increase of rye took place in the medieval period. The main reasons for introducing rye may have been changes in agricultural technology and the fact that it thrives better on poor soils than other cereals (Lange 1976a; Behre 1992). In northeastern Germany, during the Roman Iron Age, rye became firmly established as a valued crop, as evidenced by macro-botanical remains confirming its cultivation in the Roman Iron Age settlements (figure 6). In this part of Germany, rye was just as important as barley, which was also frequently cultivated (Schiemann 1957; Neef 2002; Berg-Hobohm 2004). The pollen diagrams from northeastern Germany also show a continuous presence of rye during this period (figure 2; Jahns et al 2013a; Jahns et al 2013b; Jahns 2018; Alinezhad et al 2024). However, after the Late Roman Iron Age, during the Migration Period (4^th^-6^th^ centuries CE), there is limited archaeological sites and archaeobotanical data, hindering the interpretation on the character of agriculture beyond the statement of a drop in the intensity of crop cultivation because of land abandonment. Regional pollen records from northeastern Germany from this period also indicate a gap in cereal cultivation, with a notable decline of rye (figure 2; Jahns et al 2013a; Jahns et al 2013b; Jahns 2018). According to Lityńska-Zając (1997), rye cultivation in Poland also shows a consistent and increasing presence after the Late Pre-Roman Iron Age. Rye was recorded, particularly in southern Poland, and it became significant during the Late Pre-Roman Iron Age. This trend continued into the Early Roman Iron Age, where rye was predominantly grown. Similar to Germany, pollen diagrams and archaeobotanical data from Poland, particularly in the southern and north-western regions, indicate a rise in the rye cultivation from the Late Pre-Roman Iron Age until the Late Roman Iron Age, followed by a decrease after the Late Roman Iron Age until the early Middle Ages (Lityńska-Zając 1997).

A crucial point in discussing the aforementioned interruption in cereal cultivation, particularly the decline of rye after the Late Roman Iron Age and during the Migration Period, is the presence of ergot findings and the climatic conditions during this time. The toxicity of ergot and its distinct black appearance were known, which allowed contaminated grains to be cleaned and removed. Nevertheless, archaeobotanical findings from sites dated to these periods still report ergot remains, though in low quantities. Ergot has been reported in archaeobotanical macro-remains from northern and north-eastern Germany, including sites such as Mardorf, Göritz, Schwennenz, Kablow, Groß Lübbenau, and Flögeln. In Mardorf it dates to the Pre-Roman Iron Age, but in rather low quantities. And in Flögeln it dates to the Migration Period. However, ergot findings from sites dated to the 3^rd^-5^th^ centuries CE are more frequent, such as Göritz (38 ergot finds) and Schwennenz (18 ergot finds). For Groß Lübbenau (17 ergot finds), ergot is reported during the Slavic period, but the publication does not specify whether it dates to the Early or the Late Slavic period. The storage of grains found in pits at Schwennenz was well-cleaned, with few weeds, and surprisingly it was mostly a barley stock coming along with some ergot sclerotia. Although ergot is nowadays largely associated with rye, in Schwennenz, barley is actually the only cereal species which this fungus could have infested. Behre (1992), based on the analysis of several ergot finds, concludes that this fungus existed long before the cultivation of rye and lived on other host cereals, such as barley (e.g. Kirleis 2004 for Late Bronze Age Rullstorf with emmer and barley mass finds). However, it cannot be coincidental that the first significant reports of ergotism in Europe (Alm and Elvevåg 2013) emerged alongside the spread of rye as a major crop. The expansion of its cultivation in north-eastern Germany during the 4^th^ century CE, combined with rye’s wind-pollinated nature (and not self-pollinated as the other cereals), making it more prone to ergot infestation. Thus, both its biological susceptibility and its growing role as a staple crop on poor moraine soils likely heightened the risk of ergotism under favorable climatic and environmental conditions for the spread of this fungus. While the main focus of our study is northeastern Germany we also incorporated archaeobotanical data from the old moraine in NW Germany dominated by sandy soils, and from the low mountain ranges of central Germany with loess-derived soils, to establish local and regional trends. To place these findings in a broader context, we referred to the North European Plain to the east and the Baltic region to the north, where cool-temperate climates and glacially derived soils (sandy and moraine) created comparable conditions, and where archaeobotanical and palaeoecological studies document parallel developments. For example, ergot remains have been documented in the North European Plain during the Late Pre-Roman and Roman Iron Age (Lityńska-Zając 1997), and in Scandinavia at sites such as Sorte Muld (Bornholm) and Vallhagar (Gotland) during the Migration Period (Helbæk 1957, p 272). The absence of finds from other European regions does not necessarily imply that ergot was absent there, but rather reflects the uneven availability of archaeobotanical evidence. Since rye cultivation was concentrated in particular ecological niches, such as the sandy moraine soils of northern Germany, or other areas where rye became a staple crop, these regions appear to have been especially vulnerable. Rye was particularly suited to harsh conditions, as it tolerates poor, sandy soils and cooler, wetter climates better than other cereals (Behre 1992). However, these same traits also made it more prone to ergot infestation, since rye’s wind-pollinated biology increased its susceptibility to *C. purpurea*. In contrast, areas where rye was less central, or where subsistence relied mainly on other cereals, may have been comparatively more resilient. While it remains difficult to determine whether all regions were equally affected, what is clear is that rye-dominated subsistence economies, such as those in northeastern Germany, experienced agricultural instability.

The DCAP, which was contemporaneous with the settlement’s abandonment and the reduction in cereal cultivation during the Migration Period, likely provided the climatic conditions necessary for the optimal growth of ergot. This parasitic fungus (Lev-Yadun and Halpern 2007) flourishes when cold winter is followed by an overcast wet spring and cool summer (Matossian 1989, p 13f; Shi and Yu 2022). The aforementioned regional cold, wet, and stormy conditions in regions like northeastern Germany, where rye was a staple crop during this period, likely provided the conditions for the spread of ergot, negatively impacting agricultural stability. Comparable patterns have also been suggested in other parts of Northern Europe. In line with the hypothesis that widespread epidemic ergotism may have contributed to the so-called Migration Period crisis, studies suggest it may have played a role in significant depopulation in Scandinavia and the reforestation of large agricultural areas (Gräslund and Price 2012; Alm and Elvevåg 2013; Bondeson and Bondesson 2014; Westling 2024). The first phase of abandonment in our study area during ca. 400 CE (2σ range of 330-460) to 480 CE (2σ range of 410-540) may represent a similar response, potentially linked to the agricultural instability associated with environmental stress.

The second phase of migration, as indicated by pollen results from northeastern Germany (figure 2), occurring between ca. 480 CE (2σ range of 410–540) and 550 CE (2σ range of 480–610), reflects a heightened intensity in Germanic migration. The westward push of the Huns beginning around 375 CE forced numerous Germanic groups, including the Goths, Vandals, and Alans, to cross into Roman territory, which in turn triggered large-scale demographic shifts across Europe during the Migration Period (Encyclopaedia Britannica 2025). The substantial number of people migrated from northern Germany, contributing to a marked increase in Anglo-Saxon settlers across eastern and southern England (Clarke 1985, p 40; Ejstrud et al 2008). While these socio-political factors likely played an important role in driving these migrations, environmental pressures may have further exacerbated the instability. In addition to the cool and wet conditions observed between 350–500 CE, a further climatic disturbance occurred during the LALIA, beginning with the 536 CE event. This period is associated with hemispheric cooling, environmental hazards, and social unrest across the Northern Hemisphere (Büntgen et al 2016; Toohey et al 2018; Schneeweiß 2023; van Dijk et al 2023). The darkening of the sun described by historical sources in 536 CE likely reflects the aftermath of a major volcanic eruption, resulting in several unusually cold summers and widespread crop failures (Stothers 1984; Arjava 2005; Gräslund and Price 2012).

Tree-ring evidence from Scandinavia and Central Europe, including northeastern Germany, indicates reduced summer temperatures from 536 to at least 545 CE (Gräslund and Price 2012; Arjava 2005). This aligns with our pollen-based reconstructions from lake Kleiner Tornowsee, which suggest cool and wet conditions during the late Migration Period. These environmental stressors likely contributed to the further decline in human activity and cereal cultivation already underway in the region, as shown by the noticeable decrease in anthropogenic pollen indicators and cereal-type influx (figure 2 and 4). Therefore, the 536 CE event and subsequent cooling likely intensified the existing socio-environmental instability in northeastern Germany.

## 5 Conclusion

The decline of settlement activities, cereal cultivation, and the expansion of woodlands in northeastern Germany during the 4^th^–6^th^ century CE reflect an interplay between environmental changes and human responses. Archaeological evidence from the study area shows a reduction in settlement activity from the late 4^th^ century CE onward, consistent with the migration of Germanic groups toward Roman provinces. Our pollen record shows that this demographic shift was accompanied by a marked decrease in cereal cultivation, especially rye, and an expansion of woodlands between 400 and 550 CE. Pollen-based climate reconstructions from Kleiner Tornowsee show that, following the Late Roman Warm Period, annual mean temperatures dropped by ∼2°C while annual precipitation increased, peaking at ∼850 mm. These changes coincide with a decline in *Secale* influx and a general reduction in cereal-type pollen influx. Regional stacked climate data also indicate a distinct cooling anomaly around 400 CE (1550 cal yrs BP), marked by lower summer temperatures and a concurrent increase in precipitation. Also, climate model simulations from Europe confirm pronounced cooling in northern and western regions during the 6^th^ century CE with spatially variable anomalies. These combined evidences align with the broader pattern of cold and wet conditions known as the Dark Age Cold Period (DACP) across Europe.

A notable aspect of this period is the presence of ergot (*Claviceps purpurea*) in archaeobotanical macro-remains from northern and northeastern Germany, particularly at sites dated to the Roman Iron Age/the Migration Period, 3^rd^–5^th^ centuries CE. Ergot has been reported from the Migration Period at Flögeln, where it occurs in contexts dominated by rye grains. Cereals especially rye absent from the pollen record during this time. This absence coincides with a phase of cool and wet climate conditions suggests that the cool and wet conditions of the DACP likely affected cereal cultivation by providing appropriate environmental condition for ergot spread. Under these circumstances, ergot may have negatively affected cereal productivity and food security during the 4^th^–6^th^ centuries CE.

After the decline in settlement and cereal cultivation, increased indicators of human activities become visible in the pollen record between ca. 550 CE (2σ range of 480–610) and 730 CE (2σ range of 650– 780). This phase is marked by a decrease in woodland cover and a rise in cereal pollen, especially *Secale*, along with indicators of grazing and pastoral activity, suggesting the use of woodland for pasture and the return of arable farming. A shift towards animal husbandry during this time may reflect a strategic adaptation to the preceding environmental challenges.

Archaeobotanical data from the Slavic period also show regional variation in cereal cultivation. In northeastern Germany, rye clearly became the dominant crop, which aligns with the increased *Secale* pollen in our pollen record. Additionally, the pollen-based climate reconstructions results show that conditions became better after the DACP which likely provided a more stable environment condition for agriculture.

In the above discussion, we do not argue that climatic changes or agricultural instability were the sole causes of the human migrations observed during the Migration Period. Rather, we suggest that these factors, combined with social and political dynamics, played a role. Our findings support the central hypothesis that the cool and wet conditions of the 4^th^–6^th^ centuries CE, along with the rise of rye cultivation, created favorable conditions for ergot growth and may have impacted crop yields and food security. These findings show that agricultural and demographic changes during the Migration Period were shaped by the interplay of multiple factors. Environmental stressors, combined with social and political pressures, likely had a compounding impact on settlements in northeastern Germany.

## Authorship Contribution

K. Alinezhad: research design; pollen analysis and interpretation; compiling the archaeobotanical data; writing original draft; editing. R. Geng: pollen-based climate reconstructions; review and editing. E. van Dijk: Climate Simulation model; review and editing. W. Kirleis: review and editing. M. Weinelt: Providing the Climate data; review and editing.

## Funding

The research was conducted and financed in the context of the Cluster of Excellence - EXC-2150 - ‘ROOTS – Social, Environmental, and Cultural Connectivity in Past Societies of the German Research foundation (DFG, German Research Foundation), DFG Project number 390870439.

## Statements and Declaration

Conflict of interest: The authors declare that they have no competing interest.

## Data availability statement

All data that support the findings of this study are included within the article.

## Supporting information

Archaeobotanical data

## Acknowledgements

Coring at lake Kleiner Tornowsee was conducted by O. Nelle, S. Dreibrodt, H. Usinger, W. Dörfler, M. Bahns and M. Schütz. Technical support in the laboratory was contributed by C. Floors and S. Ibens. The maps were generated with the assistance of D. Moscone using QGIS. We are grateful to all of them for their contributions.

## Notes

### Competing Interest Statement

The authors have declared no competing interest.

## References

Alinezhad K, Feeser I, Schneeweiß J, Dreibrodt S, Jahns S and Dörfler W 2024 Unravelling vegetation and human dynamics during the first millennium AD in Brandenburg, north-eastern Germany: insights from lake sediments Veget. Hist. Archaeobot. 1617–6278 10.1007/s00334-024-01032-5

Alm T and Elvevåg B 2013 Ergotism in Norway. Part 1: The symptoms and their interpretation from the late Iron Age to the seventeenth century Hist. Psychiatry 24 15–33 10.1177/0957154X11433960

Andersen S T 1979 Identification of wild grass and cereal pollen Dan. Geol. Unders. Årbog 1978 69– 92

Arjava A 2005 The mystery cloud of 536 CE in the Mediterranean sources Dumbarton Oaks Pap. 59 73–94 10.2307/4128751

Baillie M G L 1994 Dendrochronology raises questions about the nature of the AD 536 dust-veil event Holocene 4 212–217 10.1177/095968369400400211

Barclay G J and Fairweather A D 1984 Rye and ergot in the Scottish later Bronze Age Antiquity 58 126

Behre K E 1981 The interpretation of anthropogenic indicators in pollen diagrams Pollen Spores 23 225–245

Behre K E 1990 Kulturpflanzen und Unkräuter der vorrömischen Eisenzeit aus der Siedlung Rullstorf, Ldkr. Lüneburg Nachrichten aus Niedersachsens Urgeschichte 59 141–65 10.11588/nnu.1990.0.49096

Behre K E 1992 The history of rye cultivation in Europe Veget. Hist. Archaeobot. 1 141–156

Behre K-E and Kucan D 1994 Die Geschichte der Kulturlandschaft und des Ackerbaus in der Siedlungskammer Flögeln, Niedersachsen. Probleme Küstenforsch. 21 1–227.

Behre K E 2008 Landschaftsgeschichte Norddeutschlands: Umwelt und Siedlung von der Steinzeit bis zur Gegenwart. Neumünster, Wachholtz Verlag

Berg-Hobohm S 2004 Die germanische Siedlung Göritz, Lkr. Oberspreewald-Lausitz Forschungen zur Archäologie im Land Brandenburg 7 Brandenburgisches Landesamt für Denkmalpflege und Archäologisches Landesmuseum Wünsdorf

Berraies S, Rousseau E, Bérard A and Mounier J 2024 Ergot of cereals: Toxins, pathogens, and management Plant Pathol. 72 1080–1095 10.1111/ppa.13904

BGR (Bundesanstalt für Geowissenschaften und Rohstoffe) 2016 Bodenatlas Deutschland. Schweizerbart’sche Verlagsbuchhandlung, Hannover

Björck S and Clemmensen L B 2004 Aeolian sediment in raised bog deposits, Halland SW, Sweden: a new proxy record of Holocene winter storminess variation in southern Scandinavia? Holocene 14 677–688

Bjune A E, Seppä J and Birks H J B 2009 Quantitative summer-temperature reconstructions for the last 2000 years based on pollen-stratigraphical data from northern Fennoscandia J. Paleolimnol. 41 43–56 10.1007/s10933-008-9254-y

Bondeson L and Bondesson T 2014 On the mystery cloud of AD 536, a crisis in dispute and epidemic ergotism: A linking hypothesis Dan. J. Archaeol. 3 61–67 10.1080/21662282.2014.941176

Büntgen U, Myglan V S, Ljungqvist F C, McCormick M, Di Cosmo N, Sigl M, Jungclaus J, Wagner S, Krusic P J, Esper J, Kaplan J O, de Vaan M A C, Luterbacher J, Wacker L, Tegel W and Kirdyanov A V 2016 Cooling and societal change during the Late Antique Little Ice Age from 536 to around 660 AD Nat. Geosci. 9 231–236 10.1038/ngeo2652

Büntgen U, Tegel W, Nicolussi K, McCormick M, Frank D, Trouet V, Kaplan J O, Herzig F, Heussner K U, Wanner H, Luterbacher J and Esper J 2011 2500 years of European climate variability and human susceptibility Science 331 578-582 10.1126/science.1197175

Chevalier M, Davis B A S, Heiri O, Seppä H, Chase B M, Gajewski K, Lacourse T, Telford R J, Finsinger W, Guiot J, Kühl N, Maezumi S Y, Tipton J, Carter V A, Phelps L N, Dawson A, Stocker B D, Matthews-Bird F, Vallé F, Wilmshurst J M and Hickler T 2020 Pollen-based climate reconstruction techniques for late Quaternary studies Earth-Sci. Rev. 210 103384 10.1016/j.earscirev.2020.103384

Clarke H 1985 The North Sea: A highway of invasions, immigrations, and trade in the 5th to 9th centuries AD in A Bang-Andersen, B Greenhill and E H Grude ed The North Sea: A highway of economic and cultural exchange Universitetsforlaget 39–47

Clemmensen L B, Murray A, Heinemeier J, de Jong R, Madsen D B, Moska P and Nielsen L 2009 The evolution of Holocene coastal dunefields, Jutland, Denmark: A record of climate change over the past 5000 years Geomorph. 105 303-313 10.1016/j.geomorph.2008.09.024

Czerwiński S, Marcisz K, Wacnik A and Lamentowicz M 2022 Synthesis of palaeoecological data from the Polish Lowlands suggests heterogeneous patterns of old-growth forest loss after the Migration Period Sci. Rep. 12 8559 10.1038/s41598-022-12241-1

Davis B 2020 Holocene global mean surface temperature, a multi-method reconstruction approach Sci. Data 7 201 10.1038/s41597-020-0530-7

Davis M B 1963 On the theory of pollen analysis Am. J. Sci. 261 897–912 10.2475/ajs.261.10.897

Dong X, Bennion H, Battarbee R W and Sayer C D 2011 A multiproxy palaeolimnological study of climate and nutrient impacts on Esthwaite Water, England, over the past 1200 years Holocene 22 107– 118 10.1177/0959683611409778

Dreßler M, Selig U, Dörfler W, Adler S, Schubert H and Hübener T 2006 Environmental changes and the Migration Period in northern Germany as reflected in the sediments of Lake Dudinghausen Quat. Res. 66 25–37 10.1016/j.yqres.2006.02.007

Ejstrud B, Hunnicke T A, Husum C, Korre A, Maarleveld T J and Vafeiadou K 2008 The Migration Period, Southern Denmark and the North Sea. A workbook in relationship to the Gredstedbro Find. Syddansk Universitet. Marinarkæologi, Esbjerg

Esper J, Schneider L, Smerdon J E, Schöne B R and Büntgen U 2014 Northern European summer temperature variations over the Common Era from integrated tree-ring density records J. Quat. Sci. 29(5) 487–494 10.1002/jqs.2726

Faegri K and Iversen J 1989 Textbook of Pollen Analysis (4th ed.) John Wiley & Sons, Chichester

Feeser I, Dörfler W, Rösch M, Jahns S, Wolters S and Bittmann F ed 2024 Vegetationsgeschichte der Landschaften in Deutschland Springer, Berlin 10.1007/978-3-662-68936-3_1

Fang S-W, Sigl M, Toohey M, Jungclaus J, Zanchettin D and Timmreck C 2023 The role of small to moderate volcanic eruptions in the early 19th century climate Geophys. Res. Lett. 50 e2023GL105307 10.1029/2023GL105307

Fang S-W, Timmreck C, Jungclaus J, Krüger K and Schmidt H 2022 On the additivity of climate responses to the volcanic and solar forcing in the early 19th century Earth Syst. Dynam. 13 1535–1555 10.5194/esd-13-1535-2022

Geng R, Weinelt M and Zhang W 2025 Syntheses of pollen-based temperature reconstructions with respect to seasonal and spatiotemporal change in Europe Quaternary Sci. Rev. 353 109228 10.1016/j.quascirev.2025.109228

Geng R, Zhao Y, Herzschuh U, Cui Q, Zheng Z, Xiao X, Ma C and Liang C 2024 Pollen-based seasonal temperature reconstruction in Northeast China over the past 10,000 years, and its implications for understanding the Holocene Temperature Conundrum Palaeogeogr. Palaeoclimatol. Palaeoecol. 651 112391 10.1016/j.palaeo.2024.112391

Gouw-Bouman M T I J 2025 Late Holocene vegetation dynamics: degree and regional patterns of the Dark Ages woodland regeneration (AD 300–700) in the Netherlands Veget. Hist. Archaeobot. 34 29–52 10.1007/s00334-024-01000-z

Gouw-Bouman M T I J, van Asch N, Engels S and Hoek W Z 2019 Late Holocene ecological shifts and chironomid-inferred summer temperature changes reconstructed from Lake Uddelermeer, the Netherlands Palaeogeogr. Palaeoclimatol. Palaeoecol. 535 109366 10.1016/j.palaeo.2019.109366

Gräslund B and Price N 2012 Twilight of the gods? The ‘dust veil event’ of AD 536 in critical perspective Antiquity 86 428–443 10.1017/S0003598X00062862

Grauel A-L, Goudeau M-L S, De Lange G J and Bernasconi S M 2013 Climate of the past 2500 years in the Gulf of Taranto, central Mediterranean Sea: a high-resolution climate reconstruction based on δ18O and δ13C of Globigerinoides ruber (white) Holocene 23 1440–1446 10.1177/0959683613493937

Grikpėdis M and Motuzaite Matuzeviciute G 2016 The beginnings of rye (*Secale cereale*) in the East Baltics Veget. Hist. Archaeobot. 25 601–610 https://link.springer.com/article/10.1007/s00334-016-0587-6

Gunn J D 2000 The Years Without Summer: Tracing AD 536 and Its Aftermath British Archaeological Reports International (BAR) Series 872, Oxford: Archaeopress

Hamerow H, Zerl T, Stroud E A and Bogaard A 2023 The cerealisation of the Rhineland: Extensification, crop rotation, and the medieval’agricultural revolution’ in the longue durée Germania 99 157–184 10.11588/ger.2021.92156

Heiss A G, Antolín F, Bleicher N, Harb C, Jacomet S, Kühn M, Marinova E, Stika H-P and Valamoti S M 2017 State of the (t)art: Analytical approaches in the investigation of components and production traits of archaeological bread-like objects, applied to two finds from the Neolithic lakeshore settlement Parkhaus Opéra (Zürich, Switzerland) PLoS One 12 e0182401 10.1371/journal.pone.0182401

Bilbao R, Dunstone N, Menegoz M, Ortega P, Pohlmann H, Robson J I, Smith D M, Strand G, Timmreck C, Yeager S and Danabasoglu G 2020 Multi-year climate impacts of volcanic eruptions J. Geophys. Res. Atmos. 125 9 e2019JD031739 10.1029/2019JD031739

Helama S, Arppe L, Uusitalo J, Holopainen J, Mäkelä H M, Mäkinen H, Mielikäinen K, Nöjd P, Sutinen R, Taavitsainen J-P, Timonen M and Oinonen M 2018 Volcanic dust veils from sixth century tree-ring isotopes linked to reduced irradiance, primary production and human health Scientific Reports 8 1339 10.1038/s41598-018-19760-w

Helama S, Jones P D and Briffa K R 2017 Dark Ages Cold Period: A literature review and directions for future research Holocene 27 1600–1606 10.1177/0959683617693898

Helama S, Meriläinen J and Tuomenvirta H 2009 Multicentennial megadrought in northern Europe coincided with a global El Niño–Southern Oscillation drought pattern during the Medieval Climate Anomaly Geol. 37 175–178 10.1130/G25329A.1

Helama S, Stoffel M, Hall R J, Holopainen J, Jones P D, Macias Fauria M, Mielikäinen K, Timonen M, Luoto T P and Oinonen M 2021 Recurrent transitions to Little Ice Age-like climatic regimes over the Holocene Clim. Dyn. 56 3877–3893 10.1007/s00382-021-05669-0

Helama S, Timonen M, Holopainen J, Ogurtsov M G, Mielikäinen K, Eronen M, Lindholm M and Meriläinen J 2009 Summer temperature variations in Lapland during the Medieval Warm Period and the Little Ice Age relative to natural instability of thermohaline circulation on multi-decadal and multi-centennial scales J. Quat. Sci. 24 450–456 10.1002/jqs.1291

Helbæk H 1957 Bornholm plant economy in the first half of the first millennium A.D. In Klindt-Jensen O ed Bornholm i folkevandringstiden og forudsætningerne i tidlig jernalder. Nationalmuseet, København 259–277.

Herzschuh U, Bohmer T, Li C, Chevalier M, Hébert R, Dallmeyer A, Cao X, Bigelow N H, Nazarova L, Novenko E Y, Park J, Peyron O, Rudaya N A, Schlütz F, Shumilovskikh L S, Tarasov P E, Wang Y, Wen R, Xu Q and Zheng Z 2023 LegacyClimate 1.0: a dataset of pollen-based climate reconstructions from 2594 Northern Hemisphere sites covering the last 30 kyr and beyond Earth Syst. Sci. Data 15 2235–2258 10.5194/essd-15-2235-2023

Jahns S, Alsleben A, Bittmann F, Brande A, Christiansen J, Dannath Y, Effenberger H, Giesecke T, Jäger KD, Kirleis W, Klooß S, Kloss K, Kroll H, Lange E, Medović A, Neef R, Stika HP, Sudhaus D, Wiethold J, Wolters S 2018 Zur Geschichte der nacheiszeitlichen Umwelt und der Kulturpflanzen im Land Brandenburg. Beiträge zur Archäozoologie und Prähistorischen Anthropologie 11 9–35

Jahns S, Beug H J, Christiansen J, Kirleis W and Sirocko F 2013a Pollenanalytische Untersuchungen am Rudower See und Rambower Moor zur holozänen Vegetations-und Siedlungsgeschichte in der westlichen Prignitz, Brandenburg in I Heske, H J Nüsse and J Schneeweiß eds Landschaft, Besiedlung und Siedlung: Archäologische Studien im nordeuropäischen Kontext Festschrift für Karl-Heinz Willroth Wachholtz Verlag, Neumünster/Hamburg 277–293

Jahns S, Christiansen J, Kirleis W, Sudhaus D 2013b On the Holocene vegetation history of Brandenburg and Berlin in Kadrow S and Włodarczak P eds Environment and subsistence – forty years after Janusz Kruk’s “Settlement studies…”. Bonn: Dr. Rudolf Habelt GmbH 311–330Jakab G, Majkut P, Juhász I, Gulyás S, Sümegi P and Törőcsik T 2009 Palaeoclimatic signals and anthropogenic disturbances from the peatbog at Nagybárkány (North Hungary) Hydrobiologia 631 87–106 10.1007/s10750-009-9803-z

Jansma E 2020 Hydrological disasters in the NW-European lowlands during the first millennium AD: a dendrochronological reconstruction Neth. J. Geosci. 99 e11 10.1017/njg.2020.10

Jones G 2012 Weed seeds in archaeobotanical assemblages Environ. Archaeol. 17 38–53 10.1179/1461410312Z.0000000006

Juggins S 2022 rioja: Analysis of Quaternary Science Data R package version 1.0-7 https://cran.r-project.org/package=rioja

Jungclaus J, Fischer N, Haak H, Lohmann K, Marotzke J, Matei D, Mikolajewicz U, Notz D and Von Storch J 2013 Characteristics of the ocean simulations in the Max Planck Institute Ocean Model (MPIOM), the ocean component of the MPI-Earth System Model J. Adv. Model. Earth Syst. 5 422–446

Kageyama M, Albani S, Braconnot P, Harrison S P, Hopcroft P O, Ivanovic R F, Lambert F, Marti O, Peltier W R, Peterschmitt J-Y, Roche D M, Tarasov L, Zhang X, Brady E C, Haywood A M, LeGrande A N, Lunt D J, Mahowald N M, Mikolajewicz U, Nisancioglu K H, Otto-Bliesner B L, Renssen H, Tomas R A, Zhang Q, Abe-Ouchi A, Bartlein P J, Cao J, Li Q, Lohmann G, Ohgaito R, Shi X, Volodin E, Yoshida K, Zhang X and Zheng W 2017 The PMIP4 contribution to CMIP6 – Part 4: Scientific objectives and experimental design of the PMIP4-CMIP6 Last Glacial Maximum experiments and PMIP4 sensitivity experiments Geosci. Model Dev. 10 4035–4055 10.5194/gmd-10-4035-2017

Kaufman D, McKay N, Routson C et al 2020 A global database of Holocene paleotemperature records Sci. Data 7 115 10.1038/s41597-020-0445-3

Kirleis W 2004 Vegetationsgeschichtliche und archäobotanische Untersuchungen zur Landwirtschaft und Umwelt im Bereich der prähistorischen Siedlungen bei Rullstorf, Ldkr. Lüneburg, Probleme der Küstenforschung im südlichen Nordseegebiet 28 65–132

Lange E 1976a Zur Entwicklung der natürlichen und anthropogenen Vegetation in frühgeschichtlicher Zeit Feddes Repert. 87 5–30

Larsen D J, Miller G H, Geirsdóttir Á and Thordarson T 2011 A 3000-year varved record of glacier activity and climate change from the proglacial lake Hvítárvatn, Iceland Quat. Sci. Rev. 30 2715–2731 10.1016/j.quascirev.2011.05.026.

Larsen L B, Vinther B M, Briffa K R, Melvin T M, Clausen H B, Jones P D, Siggaard-Andersen M-L, Hammer C U, Eronen M, Grudd H, Gunnarson B E, Hantemirov R M, Naurzbaev M M and Nicolussi K 2008 New ice core evidence for a volcanic cause of the AD 536 dust veil Geophys. Res. Lett. 35 L04708 10.1029/2007GL032450.

Lee M R 2009 The history of ergot of rye (*Claviceps purpurea*) I: From antiquity to 1900 J. R. Coll. Physicians Edinb. 39 179–184

Legendre P and Legendre L 2012 Numerical Ecology 3rd ed Elsevier, Amsterdam ISBN 9780444538680.

Lev-Yadun S and Halpern M 2007 Ergot (Claviceps purpurea) – an aposematic fungus Symbiosis 43 105–108

Lityńska-Zając M 1997 Roślinność i gospodarka rolna w okresie rzymskim Institute of Archaeology and Ethnology, Polish Academy of Sciences, Kraków

Lukanina E, Lyubichev M, Schneeweiss J, Schultze E, Myzgin K and Shumilovskikh L 2023 Did Holocene climate drive subsistence economies in the East-European forest-steppe? Case study Omelchenki, Kharkiv region, Ukraine Quat. Sci. Rev. 305 108004 10.1016/j.quascirev.2023.108004

Luterbacher J, Werner J P, Smerdon J E, Fernández-Donado L, González-Rouco J F, Barriopedro D, Ljungqvist F C, Büntgen U, Zorita E, Wagner S, Esper J, McCarroll D, Toreti A, Frank D, Jungclaus J H, Barriendos M, Bertolin C, Bothe O, Brázdil R, Camuffo D, Dobrovolný P, Gagen M, García-Bustamante E, Ge Q, Gómez-Navarro J J, Guiot J, Hao Z, Hegerl G C, Holmgren K, Klimenko V V, Martín-Chivelet J, Pfister C, Roberts N, Schindler A, Schurer A, Solomina O, von Gunten L, Wahl E, Wanner H, Wetter O, Xoplaki E, Yuan N, Zanchettin D, Zhang H and Zerefos C 2016 European summer temperatures since Roman times Environ. Res. Lett. 11 024001 10.1088/1748-9326/11/2/024001

Matossian M K 1989 Poisons of the Past: Molds, Epidemics, and History Yale University Press, New Haven

Mauritsen T, Bader J, Becker T, Behrens J, Bittner M, Brokopf R, Brovkin V, Claussen M, Crueger T, Esch M and Fast I 2019 Developments in the MPI-M Earth System Model version 1.2 (MPI-ESM1.2) and its response to increasing CO₂ J. Adv. Model. Earth Sy. 11 998–1038

McCormick M, Büntgen U, Cane M A, Cook E R, Harper K, Huybers P, Litt T, Manning S W, Mayewski P A, More A F, Nicolussi K and Tegel W 2012 Climate change during and after the Roman Empire: Reconstructing the past from scientific and historical evidence J. Interdiscip. Hist. 43 169–220 10.1162/JINH_a_00379.

McDermott F, Mattey D P and Hawkesworth C 2001 Centennial-scale Holocene climate variability revealed by a high-resolution speleothem δ18O record from SW Ireland Science 294 1328–1331 10.1126/science.1063678

McKay N P, Kaufman D S, Arcusa S H, Kolus H R, Edge D C, Erb M P, Hancock C L, Routson C C, Żarczyński M, Marshall L P, Roberts G K and Telles F 2024 The 4.2 ka event is not remarkable in the context of Holocene climate variability Nat. Commun. 15 6555 10.1038/s41467-024-50886-w

Medović A 2004 Zum Ackerbau in der Lausitz vor 1000 Jahren. Der Massenfund verkohlten Getreides aus dem slawischen Burgwall unter dem Hof des Barockschlosses von Groß Lübbenau, Kreis Oberspreewald-Lausitz Starigard/Oldenburg– Hauptburg der Slawen in Wagrien V: Naturwissenschaftliche Beiträge, Offa-Bücher, Neumünster: Wachholtz Verlag 82 185–236

Meier M 2019 Geschichte der Völkerwanderung: Europa, Asien und Afrika vom 3. bis zum 8. Jahrhundert n. Chr. Beck C.H, München

Miedaner T and Geiger H H 2015 Biology, genetics, and management of ergot (*Claviceps spp*.) in rye, sorghum, and pearl millet Toxins 7 659–678 10.3390/toxins7030659

Neef R 2002 Ackerbau und Sammelwirtschaft. In E. Gringmuth-Dallmer and Leciejewicz E ed Forschungen zu Mensch und Umwelt im Odergebiet in ur-und frühgeschichtlicher Zeit Röm.-Ger. Forsch. 60 319–350, 401

Neukom R, Steiger N, Gómez-Navarro J J, Wang J and Werner J P 2019 No evidence for globally coherent warm and cold periods over the preindustrial Common Era Nature 571 550–554 10.1038/s41586-019-1401-2

Oosthoek J 2013 The Dust Veil Event, 536 CE. In Uekötter F (ed) What Should We Remember? A Global Poll Among Environmental Historians Glob. Environ. 11 184–214.

Orme L C, Davies S J and Duller G A T 2015 Reconstructed centennial variability of Late Holocene storminess from Cors Fochno, Wales, UK J. Quat. Sci. 30 478–488 10.1002/jqs.2792.

Peregrine P 2020 Climate and social change at the start of the Late Antique Little Ice Age Holocene 30 1–6 10.1177/0959683620941079

Pierik H J 2021 Landscape changes and human–landscape interaction during the first millennium AD in the Netherlands Neth. J. Geosci. 100 e11 10.1017/njg.2021.8

Popper V S 1988 Selecting quantitative measurements in paleoethnobotany in C A Hastorf and V S Popper ed Current Paleoethnobotany: Analytical Methods and Cultural Interpretations of Archaeological Plant Remains University of Chicago Press p 53–71

Prescott O 1813 The Natural History and Medicinal Effects of the Secale Cornutum or Ergot Dissertation Cummings and Hilliard, Boston

R Core Team 2019 R: A Language and Environment for Statistical Computing R Foundation for Statistical Computing Vienna, Austria

Ralska-Jasiewiczowa M, Miotk-Szpiganowicz G, Zachowicz J, Latałowa M and Nalepka D 2004 Carpinus betulus L. – Hornbeam in Ralska-Jasiewiczowa M, Latałowa M, Wasylikowa K, Tobolski K, Madeyska E, Wright H E Jr and Turner C eds Late Glacial and Holocene History of Vegetation in Poland based on Isopollen Maps W. Szafer Institute of Botany, Polish Academy of Science, Kraków pp 69–7

Reimann T, Tsukamoto S, Harff J, Osadczuk K and Frechen M 2011 Reconstruction of Holocene coastal foredune progradation using luminescence dating—An example from the Świna barrier (southern Baltic Sea, NW Poland) Geomorphol. 132 1–16 10.1016/j.geomorph.2011.04.017

Riechelmann D F C and Gouw-Bouman M T I J 2019 A review of climate reconstructions from terrestrial climate archives covering the first millennium AD in northwestern Europe Quat. Res. 91 111– 131 10.1017/qua.2018.84

Rørvik K-L, Grøsfjeld K and Hald M 2009 A late Holocene climate history from Malangen, a north Norwegian Fjord, based on dinocysts Nor. J. Geol. 89 135–147

Rösch M 1992 Human impact as registered in the pollen record: some results from the western Lake Constance region, Southern Germany Veget. Hist. Archaeobot. 1 101–109 10.1007/BF00206090

Rubini M, Libianchi N, Gozzi A and Zaio P 2022 The Population of the Mediterranean Basin in the First Millennium AD Migration Period Austin Anthropol. 6 1027 10.26420/austinanthropol.2022.1027

Schiemann E 1957 Die Kulturpflanzenfunde in den spätkaiserzeitlichen Speichern von Kablow bei Königs-Wusterhausen, Berliner Bl. Vor-u. Frühgesch. 5–6 100–124

Schiff P 2006 Ergot and Its Alkaloids Am. J. Pharm. Educ. 70 98 10.1016/S0002-9459(24)07817-3

Schleunes K A, Turner H A, Hamerow T S, Wallace-Hadrill J M, Barkin K, Strauss G, Kirby G H, Leyser K J, Berentsen W H, Geary P J, Duggan L G, Elkins T H, Heather P J, Sheehan J J, Bayley C C 2025 Germany Encyclopaedia Britannica https://www.britannica.com/place/Germany

Schneeweiß J 2023 Es gibt kein schlechtes Wetter – Die Folgen der Klimaanomalie von 536 n. Chr. Für die Besiedlung Norddeutschlands Hannoversches Wendland 20 49–70

Seppä H and Hicks S 2006 Integration of modern and past pollen accumulation rate (PAR) records across the arctic tree-line: A method for more precise vegetation reconstructions Quat. Sci. Rev. 25 1501–1516 10.1016/j.quascirev.2005.12.006

Seppä H, Bjune A E, Telford R J et al 2009 Last nine-thousand years of temperature variability in Northern Europe Clim. Past 5 523–535 10.5194/cp-5-523-2009

Shi H and Yu P 2022 Correlation patterns prevalence, and co-occurrence of ergot alkaloids in cool-season adapted cereal grains revealed with molecular spectroscopy and LC-MS/MS equipped HPLC system Food Chem. 393 133322 10.1016/j.foodchem.2022.133322

Silva Â, Mateus A R S, Barros S C and Silva A S 2023 Ergot Alkaloids on Cereals and Seeds: Analytical Methods, Occurrence, and Future Perspectives Molecules 28 7233 10.3390/molecules28207233

Sospedra-Alfonso R, Merryfield W J, Toohey M, Timmreck C, Vernier J P, Bethke I, Tatebe H and others 2024 Decadal prediction centers prepare for a major volcanic eruption Bull. Am. Meteorol. Soc. 105 12 E2496–E2524 10.1175/BAMS-D-23-0175.1

Stacke V, Pánek T and Sedláček J 2014 Late Holocene evolution of the Bečva River floodplain (Outer Western Carpathians, Czech Republic) Geomorphol. 206 440–451 10.1016/j.geomorph.2013.10.009

Stackebrandt W and Franke D 2015 Geologie von Brandenburg. Schweizerbart, Stuttgart Stamnes A A 2016 Effect of temperature change on Iron Age cereal production and settlement patterns in Mid-Norway in F Iversen and H Peterson ed The Agrarian Life of the North 2000 BC-AD 1000: Studies in Rural Settlement and Farming in Norway Portal Academic p 27–40. Portal Academic, Kristiansand. ISBN: 978-82-8314-099-6

Stevens B, Giorgetta M, Esch M, Mauritsen T, Crueger T, Rast S, Salzmann M, Schmidt H, Bader J, Block K, Brokopf R, Fast I, Kinne S, Kornblueh L, Lohmann U, Pincus R, Reichler T and Roeckner E 2013 Atmospheric component of the MPI-M Earth system model: ECHAM6 J. Adv. Model. Earth Syst. 5 146–172

Stothers R B 1984 Mystery cloud of AD 536 Nature 307 344–345 10.1038/307344a0

Swierczynski T, Brauer A, Lauterbach S, Dulski P, Merz B and Rohr C 2012 A 1600 yr seasonally resolved record of decadal-scale flood variability from the Austrian Pre-Alps Geol. 40 1047–1050 10.1130/G33493.1

Swindles G T, Lawson I T, Matthews I P et al 2013 Centennial-scale climate change in Ireland during the Holocene Earth-Sci. Rev. 126 300–320 10.1016/j.earscirev.2013.08.012

ter Braak C J F and Juggins S 1993 Weighted averaging partial least squares regression (WA-PLS): An improved method for reconstructing environmental variables from species assemblages Hydrobiol. 269/270 485–502 10.1007/BF00028046.

Toohey M and Sigl M 2017 Volcanic stratospheric sulfur injections and aerosol optical depth from 500 BCE to 1900 CE Earth Syst. Sci. Data 9 809–831 10.5194/essd-9-809-2017

Toohey M, Sigl M, Stoffel M et al 2016 Climatic and societal impacts of a volcanic double event at the dawn of the Middle Ages Clim. Change 136 401–412 10.1007/s10584-016-1648-7

Unger M and Lakes T 2023 Land use conflicts and synergies on agricultural land in Brandenburg, Germany Sustainability 15 4546 10.3390/su15054546

Van Dijk E, Jungclaus J, Lorenz S, Timmreck C and Krüger K 2022 Was there a volcanic-induced long-lasting cooling over the Northern Hemisphere in the mid-6th–7th century? Clim. Past 18 1601–1623 10.5194/cp-18-1601-2022

Van Dijk E, Mørkestøl Gundersen I, De Bode A et al 2024 Climatic and societal impacts in Scandinavia following the 536 and 540 CE volcanic double event Clim. Past 19 357–398 10.5194/cp-19-357-2023

Volkmann A 2014 Region im Wandel: Das 5.-6. Jahrhundert n. Chr. im inneren Barbaricum an der unteren Oder und Warthe Germania 92 133–153

Vyazov L, Myzgin K, Komar O and Rodinkova V 2024 The Western Steppe and Forest-Steppe During the Transition from the Roman Time to the Early Middle Ages in E Nikita and T Rehren ed Encyclopedia of Archaeology (2nd ed.) vol 4 Academic Press pp 870–896 10.1016/B978-0-323-90799-6.00174-9

Waltgenbach S, Riechelmann D F C, Spötl C, Jochum K P, Fohlmeister J, Schröder-Ritzrau A and Scholz D 2021 Climate variability in Central Europe during the last 2500 years reconstructed from four high-resolution multi-proxy speleothem records Geosci. 11 166 10.3390/geosciences11040166

Ward B, Pausata F S and Maher N 2021 The sensitivity of the El Niño–Southern Oscillation to volcanic aerosol spatial distribution in the MPI Grand Ensemble Earth Syst. Dynam. 12 701–716 10.5194/esd-12-701-2021

Westling S 2024 An Archaeobotanical Approach to the 6th Century Crisis: A Synthesis of Plant Macrofossil Data Exploring Agricultural Development and Resilience in Southwestern Norway Doctoral dissertation University of Stavanger, Norway ISBN: 978-82-8439-326-1

Wickham C 2009 The Inheritance of Rome: A History of Europe from 400 to 1000 Viking, LondonWood G D 2014 Tambora: The Eruption That Changed the World Princeton University Press

Wiethold J, Schäfer E and Kreuz A 2008 Archäobotanische Untersuchungen der eisenzeitlichen und kaiserzeitlichen Siedlung von Mardorf 23 in Meyer M (ed) Mardorf 23, Lkr. Marburg-Biedenkopf: Archäologische Studien zur Besiedlung des deutschen Mittelgebirgsraumes in den Jahrhunderten um Christi Geburt. Berliner Archäologische Forschungen 5, Rahden/Westf.: Verlag Marie Leidorf p 353–365

Zanchettin D, Timmreck C, Khodri M, Schmidt A, Toohey M, Abe M, Bekki S, Cole J, Fang S-W, Feng W, Hegerl G, Johnson B, Lebas N, LeGrande A N, Mann G W, Marshall L, Rieger L, Robock A, Rubinetti S, Tsigaridis K and Weierbach H 2022 Effects of forcing differences and initial conditions on inter-model agreement in the VolMIP volc-pinatubo-full experiment Geosci. Model Dev. 15 2265–2292 10.5194/gmd-15-2265-2022

